# Early enforcement of cell identity by a functional component of the terminally differentiated state

**DOI:** 10.1101/2020.01.03.894493

**Authors:** Zahra Bahrami-Nejad, Tinghuan Chen, Stefan Tholen, Zhi-Bo Zhang, Atefeh Rabiee, Michael L. Zhao, Fredric B. Kraemer, Mary N. Teruel

## Abstract

How progenitor cells can attain a distinct differentiated cell identity is a challenging problem given that critical transcription factors are often not unique to a differentiation process and the fluctuating signaling environment in which cells exist. Here we test the hypothesis that a unique differentiated cell identity can result from a core component of the differentiated state doubling up as a signaling protein that also drives differentiation. Using live single-cell imaging in the adipocyte differentiation system, we show that progenitor fat cells (preadipocytes) can only commit to terminally differentiate after upregulating FABP4, a lipid buffer that is highly enriched in mature adipocytes. Upon induction of adipogenesis, we show that after a long delay, cells first abruptly start to engage a positive feedback between CEBPA and PPARG before then engaging, after a second delay, a positive feedback between FABP4 and PPARG. These sequential positive feedbacks both need to engage in order to drive PPARG levels past the threshold for irreversible differentiation. In the last step before commitment, PPARG transcriptionally increases FABP4 expression while fatty-acid loaded FABP4 binds to and increases PPARG activity. Together, our study suggests a control principle for robust cell identity whereby a core component of the differentiated state also promotes differentiation from its own progenitor state.

**HIGHLIGHTS:**

- Fatty-acid loaded FABP4 binds to and increases PPARG expression, thereby turning on PPARG positive feedback loops that further increase PPARG expression.
- FABP4 critically controls the second phase of adipogenesis between activation of the feedback loops and reaching the threshold to differentiate.
- Only a small fraction (∼10%) of the FABP4 levels typically attained in mature fat cells is needed to commit cells to the differentiated state, thus providing an explanation for why maintenance of the mature adipocyte state is so robust.

## INTRODUCTION

Terminal cell differentiation is fundamental for developing, maintaining, and regenerating tissues in all multicellular organisms and is the process by which specialized cells such as adipocytes (fat cells), osteoblasts, neurons, and muscle cells are generated from progenitor cells [1]. However, how progenitor cells can reach a specific and unique terminally differentiated state is not well understood. In many differentiation processes, the critical transcription factors and signaling elements driving cell fate are not unique to a specific differentiation process [2–5]. Furthermore, single cell RNAseq and other single-cell technologies have provided direct evidence that cell-to-cell variability in protein expression and signaling activities can cause differentiating cells to pass through stochastic intermediate states that could misdirect cells to alternative final fates [6–9]. Given the ubiquitous use of overlapping signaling and transcription programs and the significant signaling variability between individual cells in a population, this raises the question how a differentiating cell avoids becoming lost in intermediate states and reliably finds its way to the desired specific terminal cell fate.

As one mechanism supporting terminal cell fate identity, negative feedback has been shown to play a role in regulating the differentiation decision between multiple fates whereby one fate suppress the programs that drive differentiation of the other fates [10]. However, since negative feedback mostly prevents cells which have chosen one path from differentiating into alternative cell types, additional regulatory mechanisms must exist that selectively drive cells onto a unique path and allow cells to robustly assume and maintain a specific differentiated cell identity. A number of studies focusing on directed and transdifferentiation processes showed that differentiation can proceed by multiple routes and yet converge onto similar transcriptional states [11,12], consistent with the view that terminally differentiated cell states are ‘attractor basins’ in a transcriptional and signaling differentiation landscape. The finding that the same differentiated state could be reached by passing through different intermediate states motivated us to ask the question whether a unique attractor basin requires a unique cell identify factor that would allow for the same terminally differentiated state to be reached from different intermediate states. Specifically, we considered that cells may assume a robust differentiated cell identity by using a type of positive feedback whereby a core component that is uniquely expressed and needed in the differentiated state doubles-up as a signaling co-factor that drives an irreversible step in the differentiation process.

Such a self-reinforcing mechanism that promotes robust cell identity (1) would have to control a late step before commitment to ensure that cells select the correct differentiation path and cell identity, and (2) once cells have committed, would have to robustly lock the cell into its differentiated state. To test if such a positive-feedback mechanism exists and how it may reinforce cell identity of a differentiated state, we used the adipocyte differentiation system since it is a well-characterized and experimentally accessible terminal cell differentiation process [13]. We were also intrigued by a previous observation that increased levels of fatty acids can promote differentiation of precursor cells. As a candidate for such a self-reinforcing mechanism for cell identity, the fatty acid binding protein, FAPB4, is one of the most abundant proteins in adipocytes, where it makes up between 0.5-6% of soluble protein [14]. FABP4 is normally expressed at high levels only in adipocytes [15,16] and is a critical core component of adipocytes that has important cell-internal functions such as binding to hormone sensitive lipase (HSL) and buffering lipid release [17]. However, there is also evidence that FABP4 may have an additional role in positively regulating the transition from progenitor cells into mature adipocytes [18–20]. These observations motivated our study here that FABP4 could provide such a self-reinforcement mechanism for unique cell identity. We note that FABP4 can also be released from mature adipocytes and has cell-external roles in different cell types including preadipocytes and other adipocytes where it has been shown to reduce PPARG expression as a pathological and possibly also normal regulatory function [21].

It is well-established that the expression of FABP4 is induced by PPARG, the master regulator of adipocyte differentiation [22]. However, whether FABP4 has a role in regulating PPARG activity during the differentiation process has been difficult to resolve since very little FABP4 is expressed early in adipogenesis [16,23] and that cells in the population differentiate at different times making it challenging to use traditional bulk cell assays. Also, it is not clear how FABP4 regulates PPARG and if and how FABP4 and PPARG may reinforce each other’s expression. Finally, if FABP4 is indeed uniquely important for adipogenesis, it is not clear why genetic studies in FABP4-knockout mice failed to show a suppression of adipogenesis [24].

Here, using live single-cell analysis of fluorescently-tagged endogenous PPARG and FABP4, we show that FABP4 and PPARG build up only very slowly during a first phase of the adipogenic program until they transition to a second phase marked by engagement of positive feedback between each other. This feedback starts after about 24 hours when the FABP4 level is very low and ends approximately 12 hours later when the levels of PPARG and FABP4 rapidly build up and pass a critical threshold for differentiation. We show that the feedback-activation of PPARG can be repressed by a mutant FABP4 that is deficient in binding fatty acid and that FABP4 can directly interact with PPARG. This suggests that FABP4 has - at much lower levels than seen in differentiated cells - a transport function to enhance fatty acid binding to PPARG, a mechanism which is known to increase PPARG activity [22]. Finally, we show that FABP5 may compensate for a loss of FABP4 and still allow cells to differentiate. Together, our study provides support for a likely more general model that robust cell identity can be initiated and reinforced by having a unique core component of the differentiated state double-up as a signaling factor that initiates and then reinforces the path to this unique differentiated state.

## RESULTS

### FABP4 is needed for adipogenesis in vitro and in vivo

Previous studies suggested that FABP4 and PPARG are in a positive feedback relationship [18,20,23,25] (Fig 1A). One arm of this positive feedback is well-established since there are PPARG binding sites on the FABP4 promoter, and PPARG activity has been shown to strongly upregulate FABP4 mRNA and protein levels [14]. However, the relevance of FABP4 in regulating PPARG has been controversial as FABP4 knockout mice are not defective in adipogenesis [24]. We therefore performed a series of experiments to address if and how FABP4 can regulate PPARG expression and adipogenesis. We first carried out CRISPR-mediated genome editing to completely knock out FABP4 expression in OP9 cells (Fig 1B). We then induced adipogenesis using the standard 96-hour DMI protocol and measured the percent of differentiated cells using a previously established single-cell immunocytochemistry assay based on assessing whether PPARG expression is above or below a threshold [20,23,26,27] (see Methods). As expected, control-KO OP9 cells differentiated robustly (Fig 1C). In contrast, FABP4-KO OP9 cells were strongly defective in increasing PPARG expression and differentiating, with less than 10% of cells differentiating compared to control cells. Since FABP5 has been shown to compensate for FABP4 knock-out both in vitro and in vivo [24,28,29], we also tested whether we could further reduce the amount of adipogenesis in the FABP4-KO cells by also knocking down FABP5 using siRNA. Indeed, we could reduce adipogenesis to almost nothing when we knocked down both FABP4 and FABP5 (Fig 1C).

**Figure 1:**
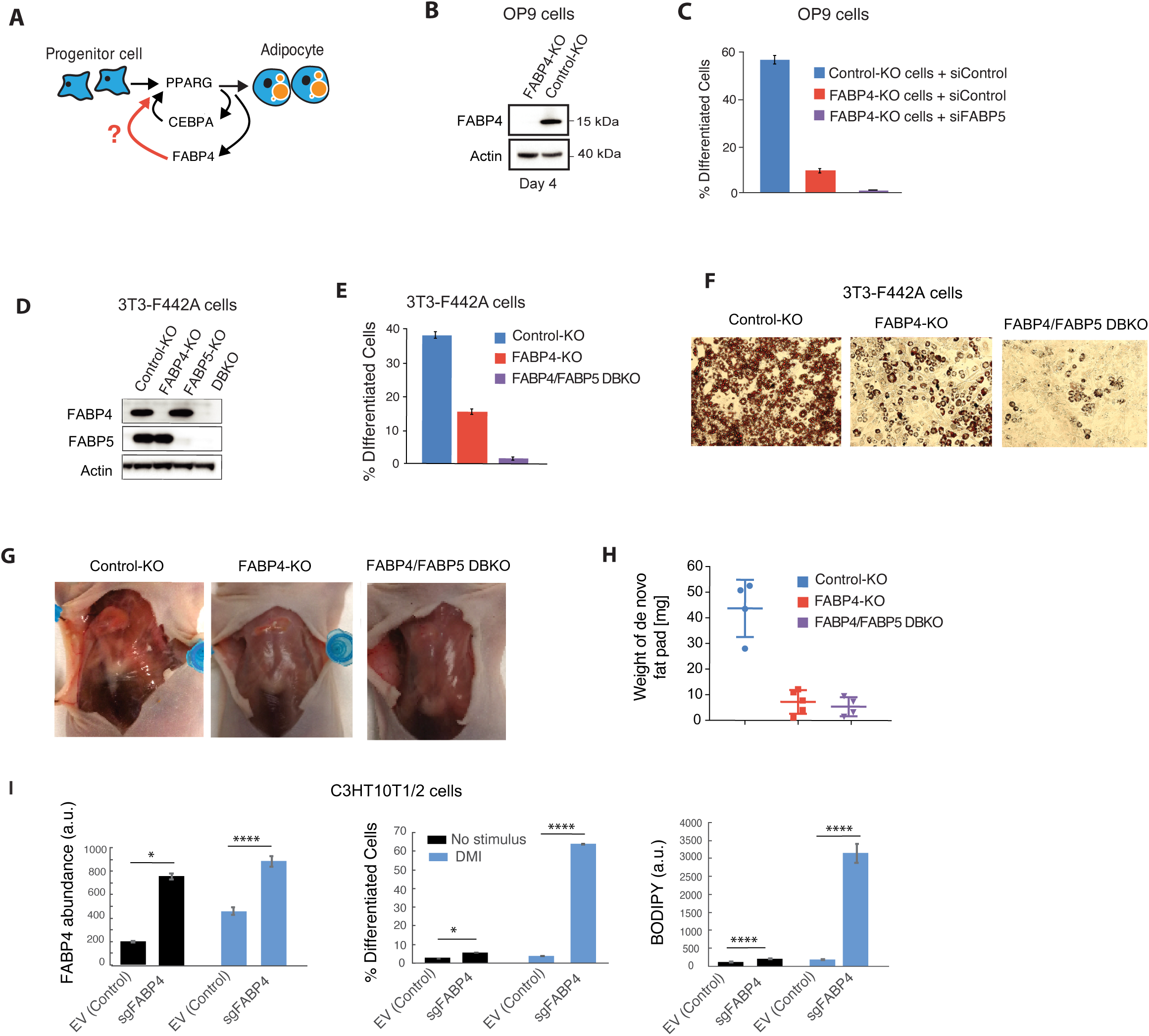
FABP4 is needed to drive adipogenesis in vitro and in vivo. (A) Schematic of proposed cell-identity positive feedback. (B-C) FABP4 was knocked out in OP9 preadipocyte cells using CRISPR-mediated genome editing. OP9 cells that were transfected and sorted at the same time but that did not harbor FABP4 knockout were used as a control. The cells were induced to differentiate by using the standard 96-hour DMI protocol (see Methods). (B) Immunoblotting of differentiated (day 4) cells. β-actin was used as a loading control. (C) PPARG expression and adipogenesis (% Differentiated Cells) were assessed by single cell immunohistochemistry as described in [20, 27]. Bar plots show mean +/– SEM of 3 technical replicates, representative of 3 independent experiments. See Supp. Fig 1A for validation of FABP5 siRNA knockdown efficiency. (D-F) 3T3-F442A cells were grown to 2 days post-confluence and were induced to differentiate by addition of insulin. Cells were assayed at day 6 post-induction. (D) Immunoblotting for FABP4 and FABP5 in FABP4-KO, FABP5-KO and FABP4/FABP5 double knockout (DBKO) 3T3-F442A cells. Wildtype 3T3-F442A cells that were transfected and sorted at the same time but that did not harbor FABP4 or FABP5 knockdown were used as a control (Control-KO). β-actin was used as a loading control. (E) The extent of adipogenesis was assessed by immunocytochemistry of PPARG. Bar plots represent mean +/- SEM from three technical replicates with approximately 1000 cells per replicate. Data is representative of three independent experiments. (F) The extent of adipogenesis was assessed by immunocytochemistry by Oil Red O staining for intracellular lipid accumulation. (G) 3T3-F442A preadipocytes subcutaneously implanted at the sternum of mice were imaged 4 weeks after implantation (H) The weight of the fat pads extracted 4 weeks after implantation. Error bars show mean + SD; unpaired t-test: *** p < 0.001. (I) C3H10T1/2 mesenchymal stem cells suitable for use to induce CRISPR-based activation of genes (CRISPRa, see Methods) were transfected with sgRNA targeting the FABP4 promoter (sgFABP4) in order to induce expression of FABP4, or with an empty vector (EV) control. Forty-eight hours later, the cells were induced to differentiate by the standard 96-hour DMI protocol as in (C). Immunocytochemistry was carried out as in (C) to measure FABP4, % Differentiated Cells, and lipid content (BODIPY). Bar plots show mean +/– SEM from 3 technical replicates and are representative of 3 independent experiments.

We further validated these findings in another commonly used preadipocyte cell system, 3T3-F442A cells [30]. We used CRISPR-mediated genome editing to knockout FABP4 or both FABP4 and FABP5 (Fig 1D). 3T3-F442A cells which had been subjected to the same FABP4 knockout protocol, but which did not harbor any FABP4 knockout were used as control cells. We plated the 3T3-F442A cells into 96-well plates and induced adipogenesis using the standard protocol in 3T3-F442A cells which is to add insulin[30]. Indeed, when FABP4 was knocked out, 3T3-F442A preadipocytes showed reduced differentiation, as measured by PPARG staining, as well as lipid accumulation, as measured by Oil Red-O staining (Fig 1E and 1F). The latter is indicative that the KO cells are not functional mature adipocytes. To test whether FABP4 was essential for adipogenesis in vivo, we used a previously-established method in which 3T3-F442A preadipocytes are subcutaneously injected into the sternum of immune-deficient mice, giving rise to fat pads resembling normal adipose tissue[31]. Since fat is not normally present at the sternum of mice, the fat pad formed at the sternum after injection of preadipocyte cells is generated by de novo adipogenesis of the injected cells[31]. We injected our preadipocyte cells into the sternum of 8-week mice, and a fat pad was allowed to form for 4 weeks. The FABP4-KO and FABP4/FABP5 double knockout (DBKO) preadipocyte cells indeed showed a defect in adipogenesis, as measured by weighing the fat that formed at the sternum (Fig 1G and 1H).

To further test for a positive feedback from FABP4 back to PPARG during adipogenesis, we next carried out overexpression experiments using CRISPR-mediated activation of FABP4 expression (CRISPRa) in C3H10T1/2 cells, a well-established mesenchymal stem cell (MSC) model which is capable of differentiating into different cell fates, including adipocytes [32]. The C3H10T1/2 cells were stably transfected with dCas9-VP64 and MS2-P65, the two core components of the CRISPRa SAM system, a second generation CRISPR-mediated activation system [33]. These C3H10T1/2-CRISPRa-SAM cells allowed us to potently activate specific genes by transfecting the cells with targeted sgRNA. To induce expression of FABP4, we transfected these cells with sgRNA targeting the promoter regions of FABP4. Two days after inducing FABP4 expression, we applied a DMI differentiation stimulus to the cells. Inducing differentiation by addition of DMI stimulus is normally not strong enough to induce adipogenesis in CH310T1/2 MSC cells. However, overexpression of FABP4 resulted in a robust increase in PPARG and adiponectin mRNA (Supp. Fig. S1), as well as in robust expression of PPARG protein and lipid accumulation (Fig 1I), compared to control cells. Since adiponectin expression and high lipid accumulation are markers of mature adipocytes, our results argue that FABP4 expression can lead to a robust increase in PPARG expression and can drive conversion of progenitor cells into mature adipocytes.

### Live-cell imaging of endogenous PPARG and FABP4 expression shows a small increase in FABP4 levels before cells reach the threshold for differentiation

Our in vitro and in vivo results (Fig 1 and Fig 2), together with previous in vitro work [18,23,25], support that PPARG and FABP4 are in a positive feedback relationship: PPARG not only increases FABP4 expression, but FABP4 can also increase PPARG expression and adipocyte differentiation. To now understand if, when, and how such a FABP4-PPARG feedback relationship functions during adipogenesis, we designed live-cell imaging experiments which allowed us to precisely determine when FABP4 and PPARG increase relative to each other in the same single cells. We used CRISPR-mediated genome editing to tag endogenous FABP4 and PPARG in mouse OP9 preadipocyte cells with orthogonal fluorescent proteins (Figs 2A, S2, S3) [20]. Using these dual-tagged citrine(YFP)-PPARG and FABP4-mKate2(RFP) cells, we carried out timecourse analysis in thousands of individual cells while applying a 48-hour long adipogenic (DMI) stimulus. As shown in Fig 2B, even though the individual time-course traces display significant cell-to-cell variability, changes in PPARG and FABP4 levels consistently show an overall similar dynamic over the four-day timecourse of adipogenesis: PPARG and FABP4 protein levels either do not increase or increase very slowly for the first 24 hours before increasing more rapidly from 24 to 48 hours, and in some cases, continuing to increase their respective abundances further for days even after the adipogenic DMI stimulus was removed at 48 hours.

**Figure 2.**
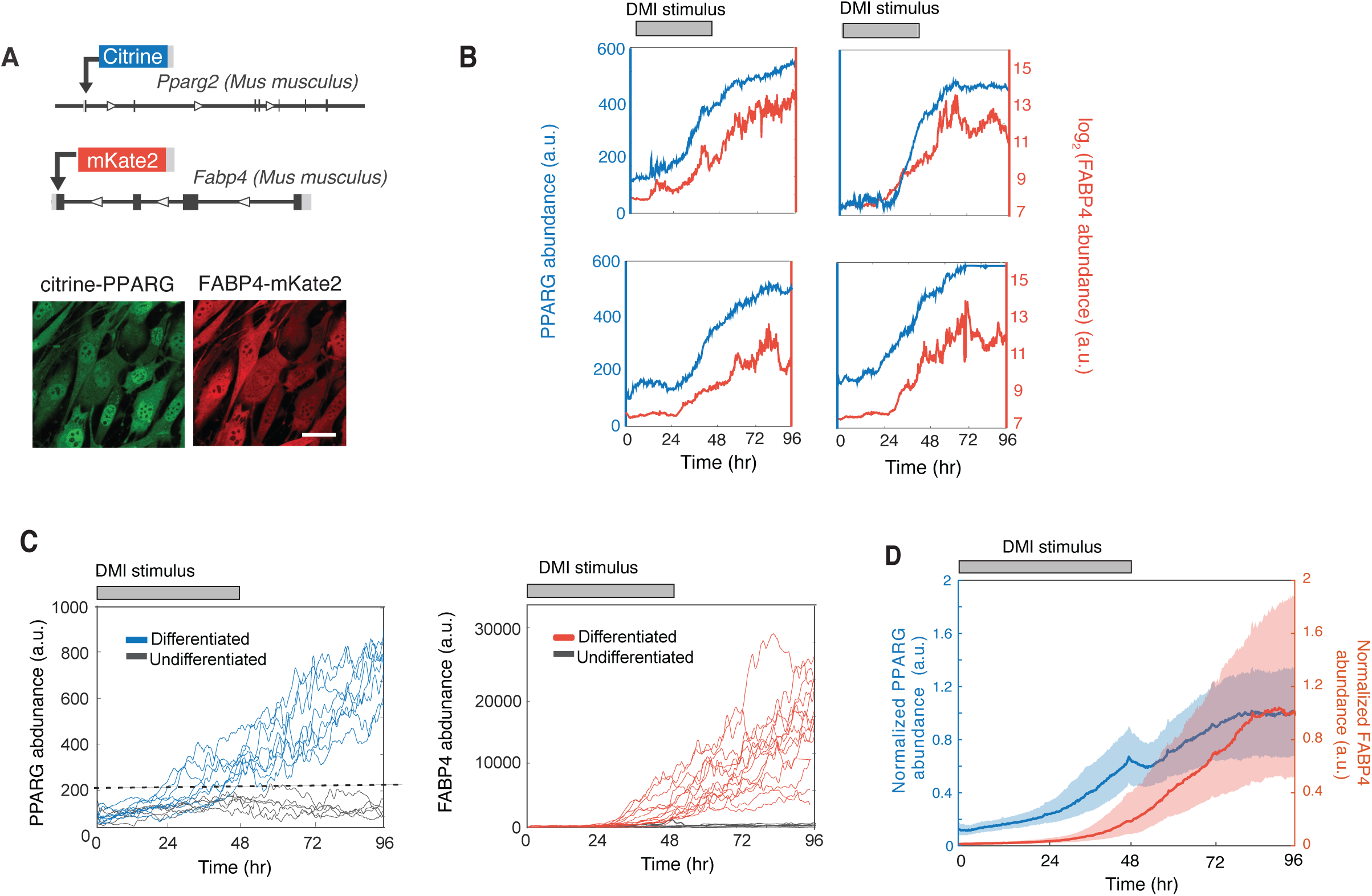
There is a delayed, small increase in FABP4 early in adipogenesis. (A) Endogenous FABP4 was tagged with mKate2 (RFP) in the citrine-PPARG cells. Images show live dual-tagged OP9 cells differentiated for 4 days into adipocytes by applying the standard adipogenic cocktail (DMI) for 2 days followed by 2 days in media with just insulin. Scale bar, 10 μM. See Supp Figs S2 and S3 for description of tagging protocol. (B) Changes in citrine-PPARG and FABP4-mKate expression measured in the same cell over the timecourse of adipogenesis. Timecourses from 4 representative cells are shown (C) Analysis of citrine-PPARG and FABP4-mKate2 timecourses in a population of differentiating cells show that there is a threshold level at which cells commit to irreversibly differentiate. (D) Normalized timecourses showing the increase in citrine-PPARG and FABP4-mKate2 levels relative to the maximal levels of the respective proteins reached in cells induced to differentiate using the standard adipogenic (DMI) protocol. Plotted lines are population median traces with shaded regions representing 25th and 75th percentiles of approximately 700 cells per condition, representative of 3 independent experiments.

In examining PPARG timecourses in the population of cells in response to the DMI stimulus, a threshold level in PPARG is apparent (marked by dotted black line in Fig 2C, left), as has been previously described [20,27]. If PPARG levels in a cell reach this threshold level, the cell will go on to differentiate even if the adipogenic stimulus is removed at 48 hours [20,27]. However, if PPARG levels in the cell do not reach this threshold level, PPARG levels in that cell will stay low or drop back down and the cell will remain undifferentiated. This analysis showed that cells that go on to differentiate according to their PPARG levels (blue traces in Fig 2C, left) also started to increase their FABP4 levels before they reach the threshold and then increase much more strongly after they reach the threshold (red traces in Fig 2C, right). Correspondingly, cells that remained undifferentiated according to their PPARG levels also did not increase their FABP4 levels (grey traces in Fig 2C).

Interestingly, even though PPARG and FABP4 protein expression dynamics look similar, the relative increase in the levels of PPARG and FABP4 is significantly different during adipogenesis. This is first indicated in the plots in Fig. 2B in which the axis for FABP4, but not PPARG, is shown logarithmically to highlight a first low range and then high range of FABP4 regulation. Over a typical 96-hour DMI differentiation timecourse, PPARG levels increase only approximately 9-fold overall compared to an approximately 120-fold increase in FABP4 levels (Fig 2D). Furthermore, in cells that eventually go on to differentiate, PPARG levels increase steadily, going up by approximately 10% of maximal by 24 hours and approximately 50% of maximal by 44 hours (Fig 2D). In striking contrast, the increase in FABP4 is markedly suppressed for the first 24 hours. Then FABP4 levels increase to only a few percent of maximal in the first 24 hours and to only 10% of maximal by 44 hours before becoming dramatically upregulated late in adipogenesis.

The small early increase in FABP4 abundance before cells reach the threshold is difficult to see without single-cell timecourse analysis. Previous studies using bulk-cell approaches such as Western blots to quantify FABP4 expression during adipogenesis lacked the sensitivity to observe the small early FABP4 increase that occurs only in the subset of cells that differentiate which is likely the reason why it has been commonly thought that FABP4 is only downstream of PPARG and is induced late in adipogenesis after the critical signaling events for differentiation have already happened [34]. Taken together, we conclude that there is a small but significant increase in FABP4 levels that occurs after a long delay after the adipogenesis program is initiated, but before cells reach the PPARG threshold at which cells commit to terminally differentiate.

### The small increase in FABP4 protein levels occurs during an intermediate step in adipogenesis and is needed to push PPARG levels up to the threshold to differentiate

When inspecting individual timecourses of FABP4 and PPARG expression in individual cells, we observed that those cells that increase PPARG early also increase FABP4 early and cells that increase PPARG at a later time also increase FABP4 later. This correlation can best be seen by dividing timecourses from a typical differentiation experiment into 4 bins and comparing how the average PPARG and FABP4 levels in the corresponding bins change (Figs 3A and 3B). Indeed, cells that increase PPARG early (red), near the median (blue), or late (green), as well as cells that do not differentiate (grey), have corresponding changes in FABP4 levels. When focusing on the moment when PPARG levels start to increase (Fig 3B), it is apparent that there is a rapid change in the slopes of both the PPARG and FABP4 timecourses between 18 and 36 hours after adipogenic stimulus is added. This change in the rate of increase of PPARG levels is apparent even in individual PPARG traces (Fig 3C). Such a delayed and abrupt change in slope is a hallmark of positive feedback that may involve both FABP4 and PPARG, as well as other established positive feedback partners of PPARG such as CEBPA [23,35], enhancing the activity and expression of PPARG (Fig 3D). We thus refer to this distinct change in the rate of PPARG expression between approximately 18 to 36 hours, which can perhaps best be seen in a slope analysis (Fig. 3E), as the timepoint when the positive feedbacks to PPARG start to engage. We had previously observed that the feedback engagement point occurs before cells reach the threshold to differentiate [20]. To validate this here, we carried out siRNA experiments. Indeed, knockdown of the two PPARG feedback partners, CEBPA and FABP4, resulted in a failure to reach the PPARG threshold (Fig 3F). Interestingly we found that the CEBPA feedback engages before the FABP4 feedback (Fig 3F).

**Figure 3.**
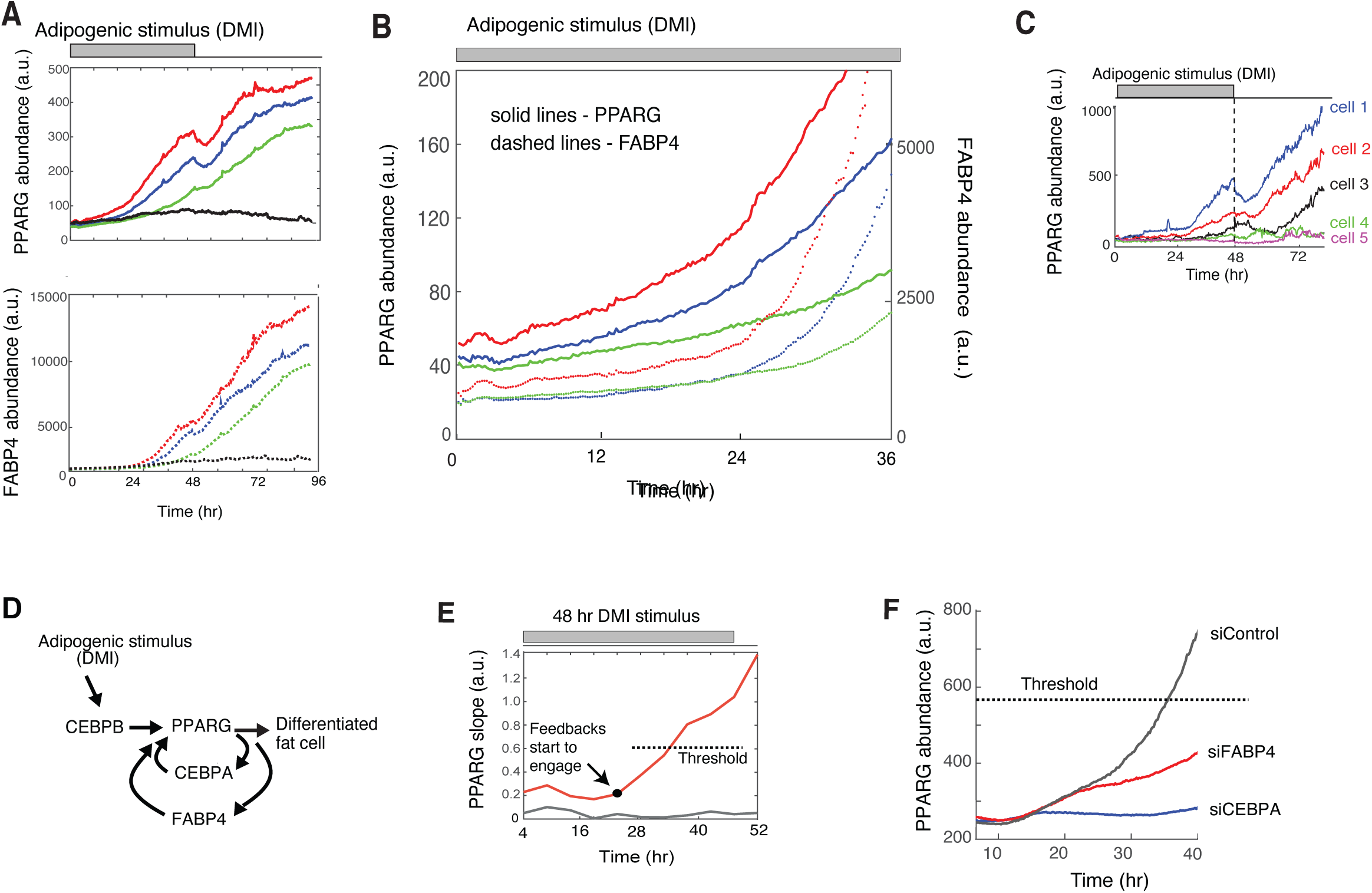
The early, small increase in FABP4 is needed to push PPARG levels to the threshold to differentiate. (A) Timecourses of 494 dual-tagged cells were binned according to their PPARG level at 48 hours, and the traces in each bin were averaged. Plot shows representative averaged timecourses to show a range of differentiation outcomes. The data shown is representative of 5 independent experiments. (B) Zoomed-in region of (A) showing a direct comparison of the gradual increases in PPARG and FABP4 before both get self-amplified when cells reach the engagement point (C) Examples of single-cell timecourses of PPARG expression in citrine-PPARG OP9 cells stimulated to differentiation using the standard 96-hour DMI protocol. (D) Schematic of the relationship between CEBPB, CEBPA, FABP4, and PPARG to control adipogenesis. (E) Red line shows the average slope of ∼700 PPARG timecourses in a typical DMI-induced differentiation experiment. The slope of PPARG at each timepoint was calculated by using a linear fit to 8-hour segments of the PPARG abundance trajectory (+/- 4 hours). The timepoint when the feedbacks start to engage is reflected in a change in PPARG slope and occurs many hours before a cell will cross the PPARG threshold level at which differentiation is irreversibly triggered. Data is representative of five independent experiments. (F) Citrine-PPARG OP9 cells transfected with siRNA targeting CEBPA, FABP4, or scrambled control and stimulated to differentiate using the standard 96-hour DMI protocol. Plotted lines are population median traces with shaded regions representing 25th and 75th percentiles of approximately 700 cells per condition, representative of 6 independent experiments.

To better determine when the different regulators of adipogenesis become important to drive the terminal differentiation process, we carried out more in-depth analysis of how these proteins influence PPARG increases during adipogenesis. CEBPB is a CEBPA homolog that is believed to function early in adipogenesis (Fig 3D) [36]. We thus considered that CEBPB may be required first and that the two positive feedbacks centered on FABP4 and CEBPA may then act either in parallel or sequentially during the feedback engagement point following CEBPB activation (Fig 3D) [20]. Indeed, as shown in Fig 4A, knockdown of CEBPB suppresses the initial gradual PPARG increase already at approximately 4 hours after adipogenesis, supporting a critical role of CEBPB in mediating the initial slow gradual increase in PPARG that occurs before the positive feedbacks engage. In contrast, knockdown of CEBPA suppresses the normal PPARG increase only at approximately 17 hours after differentiation is induced (Figs 3F and 4A). Strikingly, knockdown of FABP4 suppresses the normal PPARG increase much later, at approximately 27 hours after stimulation (Figs 3F and 4A), arguing that the CEBPA-PPARG and FABP4-PPARG feedback loops act sequentially to drive the increase in PPARG in an intermediate phase of the differentiation process before the threshold is reached at approximately 32 hours after DMI stimulation (Fig 4B). As shown in the timecourses in Figs 2C and 4B, cells reach the threshold at different times, but the average threshold time converges to one value (histogram, Fig 4B).

**Figure 4.**
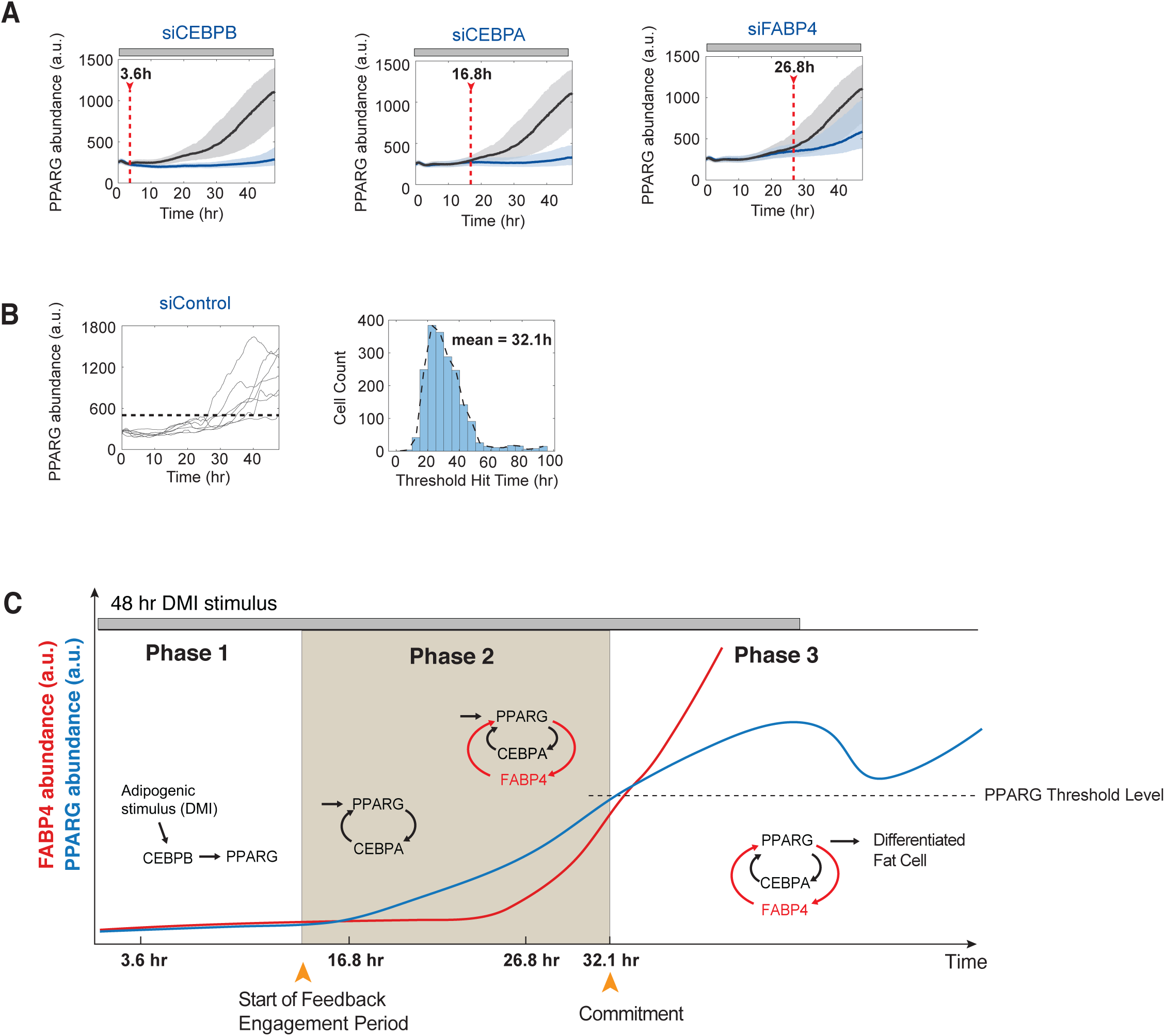
During adipogenesis, the PPARG-FABP4 feedback engages after the PPARG-CEBPA feedback engages. (A) Plots show timecourses in citrine-PPARG OP9 cells transfected with siRNA targeting CEBPB, CEBPA, FABP4 (blue lines and shades) or scrambled control (gray lines and shades) and stimulated to differentiate using the standard 48-hour DMI protocol. Plotted lines are population median traces with shaded regions representing 25th and 75th percentiles of approximately 700 cells per condition, representative of 6 independent experiments. For each knockdown timecourse, the time point at which the targeted protein becomes engaged in adipogenesis is estimated by determining when the traces of knockdown cells start to diverge from that of control cells (see details in Supp. Fig. S5). (B) Random examples of single-cell timecourses of PPARG expression in scrambled control cells from (A) show that the timing to reach PPARG threshold varied across the population (left panel). However, the histogram of the threshold hit times from single cells is strongly unimodel with a mean value of 32.1 hour (right panel). (C) Schematic illustrating the three phases of adipogenesis. FABP4 is not active early and only starts to become active later in differentiation to control the critical middle phase of differentiation (Phase 2) during which cells commit irreversibly to differentiate. During Phase 1, PPARG slowly increases to the feedback engagement point, primarily driven by the external DMI stimulus. During Phase 2, both external stimulation and internal self-amplification are needed to increase PPARG levels up to the threshold. In Phase 3, which begins after the threshold is reached, the cell trajectory becomes independent of input stimulus, and a further increase can be driven by internal self-amplification alone until the terminal differentiation state is reached. The observed drop in PPARG level is due to loss of the input stimulus before PPARG levels once again increase due to positive feedback to PPARG.

Our results are consistent with the schematic shown in Fig 4C: CEBPB drives the initial slow gradual increase in PPARG during an initial Phase 1, which is then followed by an intermediate Phase 2 of adipogenesis when a first positive feedback between PPARG and CEBPA engages. After a delay, this first positive feedback is further amplified by a second positive feedback between PPARG and FABP4 that engages just before cells reach the threshold for differentiation. Once cells reach the PPARG threshold, high PPARG levels become self-sustaining, being driven by positive feedback independently of input stimulus. In this maintenance phase (Phase 3), the positive feedbacks stay active and cause further increases in PPARG levels, perhaps to lock cells more strongly in the differentiated state. In this Phase 3 of the differentiation process, FABP4 levels also dramatically increase to much higher levels as shown in the timecourses in Fig 3. Of note, the PPARG threshold, which marks the transition between Phase 2 and Phase 3, cannot be seen just by inspecting the single-cell or averaged traces and needs to be identified by using live-cell imaging to measure PPARG levels before removing the external differentiation stimulus and then continuing to track each individual cell to know their final differentiation state days later [20,37]. Taken together, our results support that the intermediate phase of adipogenesis, Phase 2, is driven by engagement of first CEBPA and then FABP4 to trigger sequential positive feedbacks that drive PPARG to the threshold necessary for cells to become irreversibly differentiated.

### FABP4 is needed to transport lipid ligand to PPARG in order for preadipocytes to differentiate

Previous studies proposed that FABP4 may have a role to deliver fatty-acid ligands from the cytosol to the nuclear receptor PPARG [18], thereby enhancing the transcriptional activity of PPARG. Since such a role of FABP4 has been controversial, we took advantage of our knockout cell line to test whether the transcriptional activity of PPARG is indeed enhanced by FABP4 being able to transport lipid ligand to PPARG. To test for the importance of fatty acid binding, we made use of an R126Q mutation in FABP4 that results in a 30-50-fold reduction in the binding affinity for long chain fatty acids [17,38]. We used the FABP4-KO OP9 preadipocyte cells we had previously generated (described in Fig 1D) and made stable cell lines expressing doxycycline-inducible YFP-FABP4(WT), YFP-FABP4(R126Q) non-fatty acid binding mutant, or YFP(control) (Fig 5A). Inducing expression of wildtype FABP4 in the FABP4-KO cells significantly increased adipogenesis over control cells (Fig 5B). In contrast, FABP4-KO cells in which FABP4-R126Q, a non-fatty acid binding mutant of FABP4, was expressed resulted in minimal differentiation with PPARG levels dropping well below control levels (Fig 5B). This result suggests that the mutant FABP4 can suppress PPARG in a dominant-negative manner.

**Figure 5.**
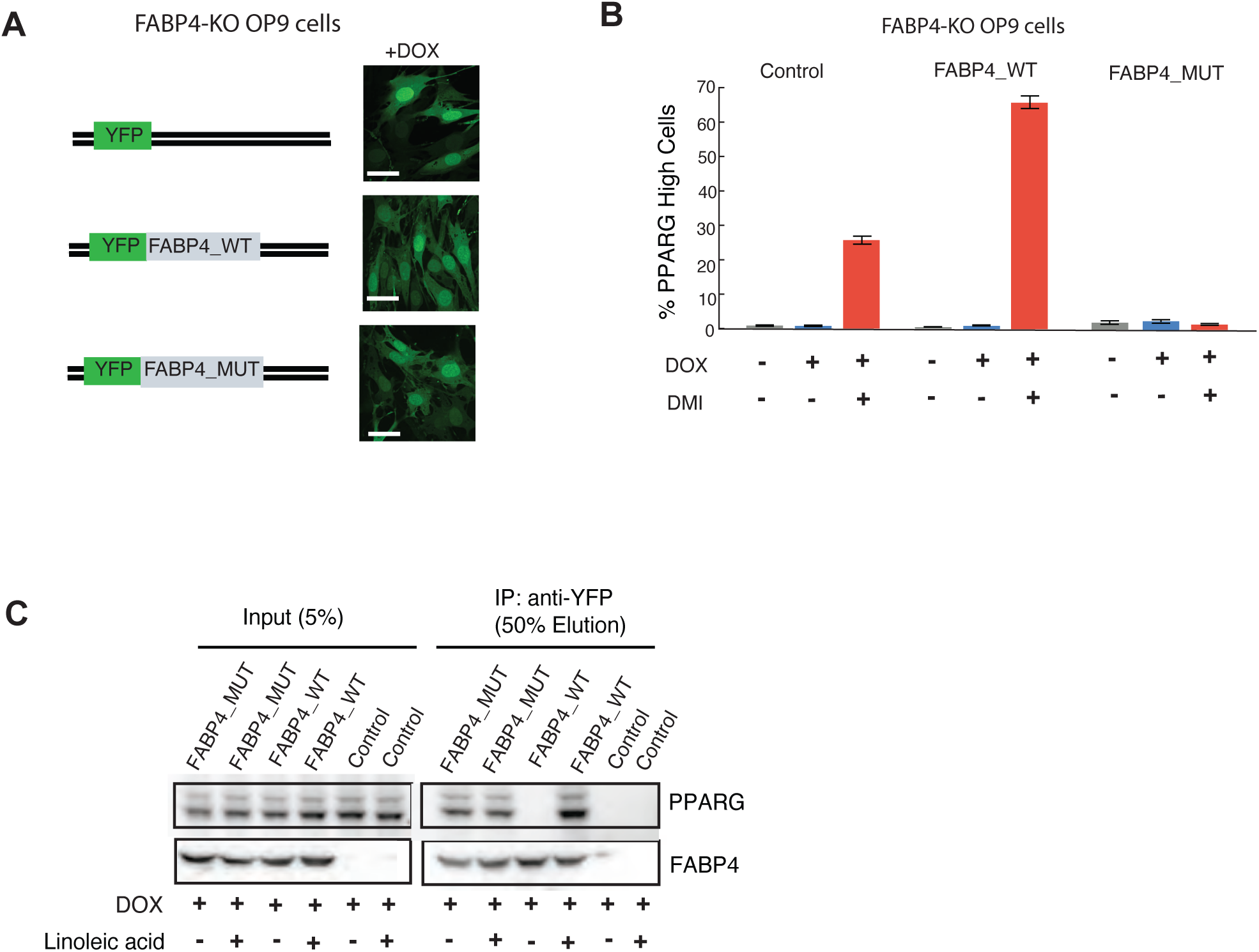
The lipid-binding activity of FABP4 is needed to increase PPARG expression and induce adipogenesis. (A) Generation of stable cell lines in which YFP-FABP4(WT), non fatty-acid binding mutant YFP-FABP4(R126Q), or YFP(control) can be overexpressed in a doxycycline-inducible manner in the FABP4-KO OP9 cell line. (B) Overexpression of YFP(control) or YFP-FABP4(WT) in OP9 cells resulted in maximal differentiation. In contrast, overexpression of YFP-FABP4(R126Q), a non-fatty acid binding mutant of FABP4, resulted in reduced differentiation compared to control. Overexpression of YFP-FABP4(WT) in FABP4-KO cells rescued the differentiation capacity. In contrast, overexpression of YFP(control) or YFP-FABP4(R126Q) resulted in no differentiation. (C) To assess PPARG interaction with FABP4, co-IPs on total cell lysate were performed using GFP nanobody or IgG(control) antibodies in FABP4-KO cells with overexpressin YFP-FABP4(WT) or YFP-FABP4(R126Q) mutant. At t=0 hours, cells were treated with both linoleic acid (100 μM) and doxycycline (1ug/ml). Co-IP’s on the total cell lysates were performed at day two (t=48 hours). The Western blot for FABP4 and PPARG is representative of 2 independent experiments.

To test whether FABP4 and PPARG directly interact, we added doxycycline to cells described in Fig 5A to overexpress YFP-FABP4, YFP-FABP4-mutant, or YFP(control) and performed a co-immunoprecipitation (coIP) assay using a GFP-nanobody (Fig 5C). We found that FABP4 and PPARG indeed interact in a lipid-dependent manner, suggesting that lipid-loaded FABP4 can directly transfer lipid to PPARG. Interestingly, we found that the fatty-acid binding deficient mutant-FABP4 is bound to PPARG regardless of whether fatty acid was present or not (Fig 5C), explaining the observed dominant negative effect of the FABP4 mutant which likely acts by preventing endogenous FABPs from transferring fatty acid to and activating PPARG. Together these results support the hypothesis that FABP4 can function as a fatty acid transfer protein that increases PPARG activity to commit cells to terminally differentiate.

### Rosiglitazone is a small molecule, high-affinity activator of PPARG that can bypass the need for FABP4 to transport lipid to PPARG

Given the results thus far that FABP4, or FABP5 in a compensatory role, is needed to increase PPARG expression to high enough levels to switch progenitor cells into adipocytes, there is a puzzling finding in the literature that mouse embryonic fibroblasts (MEFs) from FABP4 and FABP5 double-knockout mice can still differentiate into adipocytes[39]. We tested whether the contradiction could be due to the fact that rosiglitazone, a small molecule activator of PPARG, had been added to the standard cocktail used to induce adipogenesis. We were indeed able to reproduce this result by also adding rosiglitazone to the adipogenic cocktail. In Figs 1C and 1E, we had shown that knockout of FABP4 alone, or together with knockdown or knockout of FABP5, resulted in a dramatic reduction in PPARG expression and differentiation in response to an adipogenic stimulus. However, when we added rosiglitazone along with the adipogenic stimulus, the differentiation capacity of the FABP4-KO cells was restored in the FABP4-KO OP9 and 3T3-F442A cells (Figs 6A, 6B, S1B, and S1C). Approximately 50% of the differentiation capacity was restored in the FABP5-KO and double knockout 3T3-F442A cells, and an even higher fraction of differentiation was restored for the FABP5-KD and double knockout OP9 cell models.

**Figure 6.**
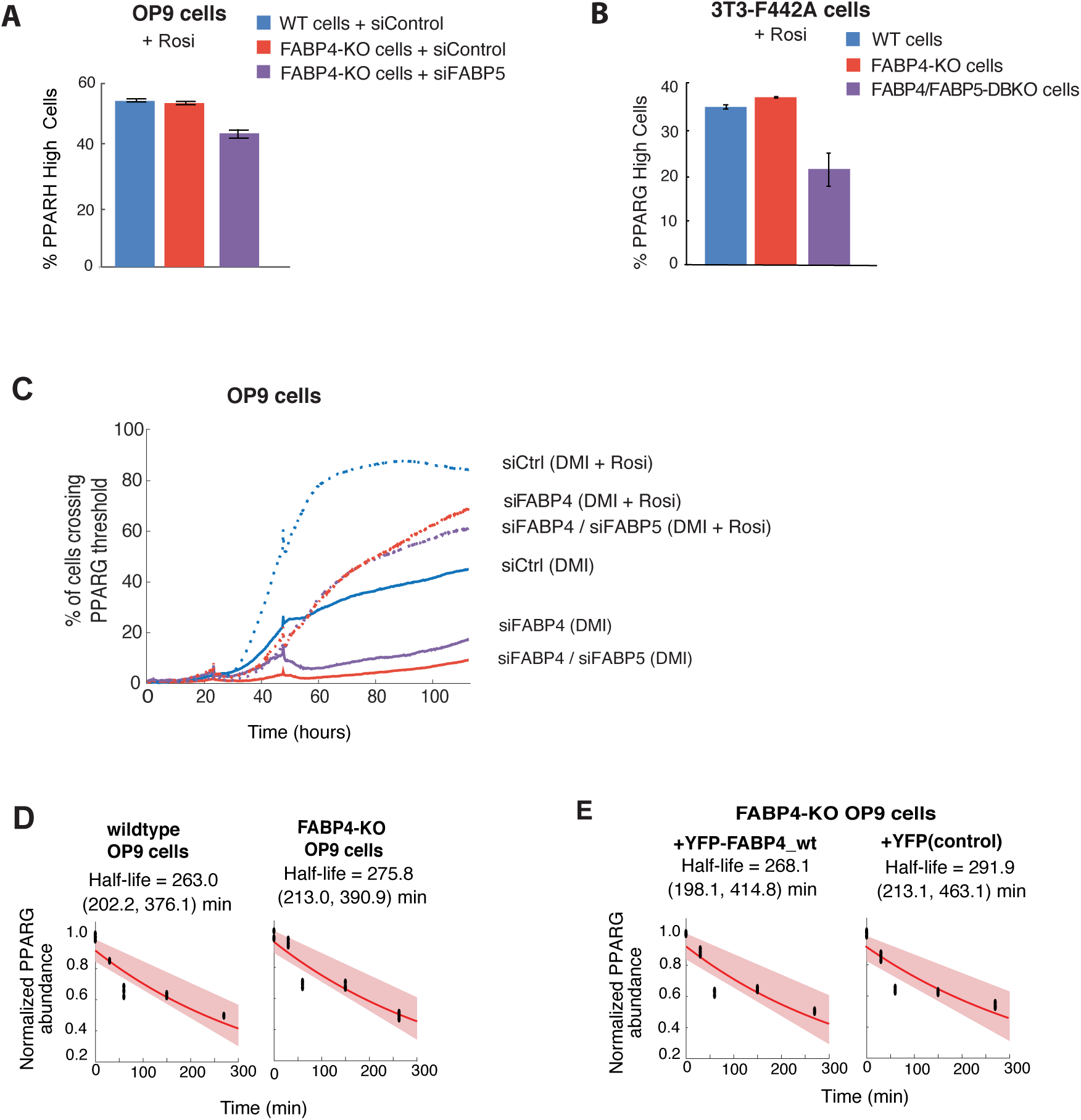
Rosiglitazone, a PPARG agonist which binds to the same binding pocket in PPARG as fatty acid, bypasses the need for FABP4 upregulating PPARG expression during adipogenesis. (A) Addition of Rosiglitazone (1 μM) to the differentiation stimulus (DMI) rescues the loss of adipogenesis shown in Fig.1E in FABP4-KO OP9 cells and in FABP4-KO OP9 cells transfected with FABP5 siRNA. Percent of differentiated cells was quantitated as in Figs 1B and 1C. Images are shown in Supp Fig S1. (B) The addition of Rosiglitazone (1 μM) to the differentiation media (insulin added to the growth media) rescues the loss of adipogenesis in FABP4-KO, FABP5-KO, and DBKO 3T3-F442A cells shown in Figs 1G-1H. Percent of differentiated cells was quantitated as in Fig 1G. Images are shown in Supp Fig S1. (C) Citrine-PPARG cells were transfected with FABP4 and FABP5-targeted or control (YFP) siRNA and were imaged while being induced to differentiate by the standard 96-hour DMI protocol with and without Rosiglitazone added to the DMI stimulus. Knockdown of FABP4 and FABP5 results in loss of differentiation. However, this phenotype was partially rescued by addition of Rosiglitazone. (D-E) Wildtype OP9 cells FABP4KO-YFP-FABP4 or FABP4KO-YFP(control) cells were stimulated with lineolic acid for 48 hr to induce FABP4 to PPARG binding and were then treated with cycloheximide and fixed at the indicated time points. Immunocytochemistry was performed to measure PPARG protein levels. The shaded region represents the 95th confidence bounds on the fitted coefficients. Values enclosed in parentheses show lower and upper bounds on the estimated half-life derived from the 95% confidence bounds.

**Figure 7.**
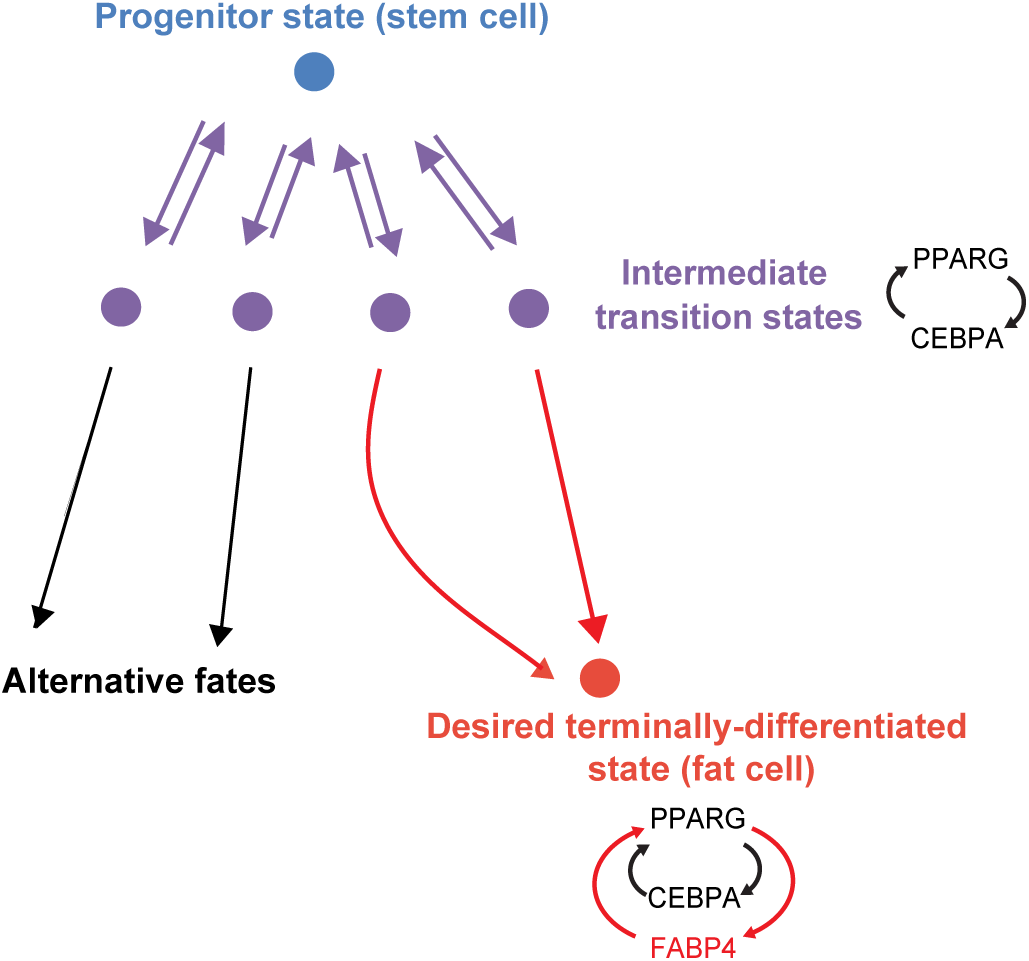
Model for how FABP4 functions as an attractor molecule for the terminally differentiated state.

A live-cell analysis of rosiglitazone addition in cells with endogenously tagged PPARG provides mechanistic insights how rosiglitazone acts (Fig 6C). In DMI-stimulated control cells, addition of rosiglitazone has little effect early in adipogenesis but then causes a much more rapid increase in PPARG levels when the feedback engagement point is reached after 24-36 hours. For the same conditions with and without rosiglitazone, FABP4 or FABP4/5 knockdown suppresses the rapid increase in control cells. However, rosiglitazone partially restores the fast increase of PPARG in both knockdown cells. Our results can be interpreted that rosiglitazone, which is soluble in water at low concentrations and has a very high affinity for PPARG [40], can directly access the activating binding pocket of PPARG without needing a cell-internal lipid transport mechanism provided by FABP4 or FABP5. In contrast, fatty acids, which have a much lower solubility in water, require a lipid transport mechanism to reach PPARG and this transport function requires either FABP4 or FABP5 in these cell systems.

One question is how can FABP4, which is not a transcription factor, increase PPARG expression? Since FABP4 can directly bind to PPARG in the presence of lipid (Figure 5D), it is conceivable that the binding may increase PPARG expression indirectly by stabilizing PPARG. To test for such a potential stabilization, we measured degradation rates of PPARG in two conditions that compared when lipid-loaded FABP4 was bound to PPARG versus when no FABP4 was bound to PPARG. In both conditions linoleic acid was added to cells for 2 days to induce FABP4 binding to PPARG (Fig 5D). In condition 1 (Fig 6D), we compared wildtype OP9 preadipocyte cells which expressed FABP4 versus to OP9 cells in which FABP4 had been knocked out (FABP4-KO cells from Fig. 1D). In condition 2 (Fig 6E), we compared FABP4-KO cells which either overexpressed YFP-FABP4 or a YFP(control). In all cases, there was no significant difference in PPARG degradation rate, suggesting that binding of lipid-loaded FABP4 to PPARG does not stabilize PPARG. Together, these results provide support for the model that lipid-loaded FABP4 increases PPARG expression by bringing lipid to PPARG to increase PPARG activity and not by stabilizing PPARG.

## DISCUSSION

Our study focuses on a general question how cells ensure that they assume and maintain a specific terminally differentiated state given that differentiating cells typically have multiple cell-fate options and there is also cellular plasticity with cells being able to transition between different reversible progenitor states. Our study considered that one solution to the cell identity problem is to reinforce the differentiation path to a specific differentiated state by using a unique core component of the differentiated state also in a separate regulatory capacity to both control a key step in the differentiation process and to then maintain the differentiated state. Here we tested whether FABP4, one of the most abundant proteins in mature adipocytes that regulates lipid flux, could function as such a cell identity factor in adipogenesis.

Our rationale for focusing on FABP4 was previous evidence that FABP4 may regulate PPARG expression and/or activity [18,20,23,25] and that FABP4 is highly expressed specifically in adipocytes [15]. We investigated the role of FABP4 by using simultaneous single-cell analysis of endogenously tagged FABP4 and PPARG to test if and when FABP4 and PPARG regulate each other. We used this type of analysis in order to reconcile the seemingly conflicting results in the literature that we thought might be tied to a high cell-to-cell variability in the dynamics of FABP4 expression and the possibility that low levels of FABP4 might be sufficient to activate PPARG. By tracking cells and monitoring expression of PPARG in individual cells over the timecourse of DMI-induced adipogenesis, we showed that PPARG expression during adipogenesis can be divided into three sequential Phases with DMI mediating a linear increase in PPARG in Phase 1, positive feedback starting to engage in Phase 2 and ending with PPARG levels reaching a critical threshold where self-amplification takes over and the external input stimulus is not needed any more. Markedly, we showed that this Phase 2 is started by a first positive feedback between PPARG and CEBPA and is about 8 hours later amplified by a second positive feedback between PPARG and FABP4. In Phase 3, this same positive feedback between FABP4 and PPARG helps maintain the differentiated adipocyte state.

We showed that FABP4 is a necessary co-factor both in OP9 and 3T3-F442A preadipocyte cells to accelerate the build-up of PPARG in Phase 2 to the threshold where differentiation is triggered. However, as shown in Fig. 3D, only low levels of FABP4 - less than 10% of the maximally expressed FABP4 - are needed for this role of FABP4 in increasing PPARG activity which ensures first robust induction and then robust maintenance of the terminally differentiated state. Taken together, our study suggests that FABP4 has at least two roles in adipocytes: to transport and buffer fatty acids in mature cells during lipolysis, and at very low levels, to act in adipocyte precursor cells as a signaling mediator that transfers fatty acids to PPARG and increase its activity to promote differentiation.

Interestingly we found that the FABP4-PPARG feedback engages many hours after the CEBPA-PPARG positive feedback, offering a possible explanation of how to selectively achieve the mature adipocyte fate despite that the PPARG and CCAAT/enhancer binding protein transcription factors are common in many cell types [2,41–44]. It is possible that in the many cell types that express PPARG and CEBPA, PPARG levels can only reach an intermediate level when the PPARG-CEBPA feedback loop engages (Figs 3F and 7). Only when FABP4-PPARG positive feedback sequentially engages can PPARG and FABP4 levels rise so strongly and rapidly that the differentiating cell can selectively reach the final adipocyte differentiated state. Our findings may have implications for treating insulin resistance, diabetes, and other metabolic diseases for which defective adipogenesis and lack of functioning fat cells to properly store and release lipids is a hallmark. Drugs such as rosiglitazone that potently and directly activate PPARG are believed to treat insulin resistance in part by promoting adipogenesis and fat cell function, but they have side effects since PPARG is critical in other differentiation and regulatory processes in different tissues. Our finding that adipogenesis is selectively driven by a druggable and late-acting adipocyte specific factor FABP4 may ultimately end up being a useful selective strategy to therapeutically control adipogenesis.

Our study further addressed the question of how FABP4 can regulate PPARG since previous studies have yielded contradictory results. Some studies suggested that FABP4 upregulates PPARG expression and activity in adipocytes [18,23,25] whereas others suggested that FABP4 suppresses PPARG expression and activity [21,45]. We showed that knocking down or knocking out both FABP4 and FABP5 in OP9 cells and 3t3-F442A results in a significant suppression of adipogenesis. This first appeared to contradict earlier results which showed that WT cells, FABP4/FABP5 KO cells, FABP4/FABP5 KO cells can still effectively differentiate when rosiglitazone was added along with the DMI. Our study provides an explanation for this result, since we found that these double knockout cells fail to respond to DMI but differentiate in response to the PPARG activator rosiglitazone which was included in the previous experiments. We show that FABP4 can directly interact with PPARG and that a mutant deficient in fatty acid binding has increased PPARG affinity without bound fatty acid. Together, these results suggest that rosiglitazone can bind directly to the lipid-binding pocket of PPARG and may therefore bypass the need for FABP4 to bring fatty acids to PPARG and activate PPARG. Such a mechanism is plausible as fatty acids have low water solubility while rosiglitazone was developed as a partially water-soluble compound. Together with our knockout, mutant and rescue analysis, we also used live-cell imaging and demonstrated a rapid increase of endogenously tagged PPARG activity mediated by rosiglitazone. Thus, by mediating direct activation, rosiglitazone can bypass the need for FABP4. We therefore conclude that FABP4 has a dual role in adipocytes, as an essential lipid buffer and transporter that promotes lipid metabolism in the differentiated state, but also as an essential co-factor for terminal cell differentiation that drives precursor cells towards a specific commitment step and then helps maintain adipocyte cell identity.

## AUTHOR CONTRIBUTIONS

Z.B., S.T., A.R., M.L.Z., and M.N.T. designed and carried out experiments and analyzed data. T.C. and Z.Z. analyzed data. F.B.K. designed experiments and contributed to the discussion. Z.B., T.C., and M.N.T. wrote the manuscript with input from all authors.

## ACKNOWLEDGEMENTS

This work was supported by National Institutes of Health RO1-DK101743, RO1-DK106241, P50-GM107615, P30 DK116074, a Stanford BioX Seed Grant, and a Stanford Diabetes Research Center Seed Grant (to M.N.T.), 1F31DK112570-01A1 (to M.L.Z.), DFG Research Fellowship TH2156/1-1 (to S.T.), NovoNordisk (to A.R.), MOST Fellowship and Stanford Deans Fellowships (to T.C.), and by the Department of Veterans Affairs (Office of Research and Development, Medical Research Service) to F.B.K. We thank Tobias Meyer (Stanford University) and members of the Teruel Lab for discussions and critical reading of the manuscript.

## SUPPLEMENTARY FIGURE LEGENDS

**Figure S1. Additional experiments supporting that FABP4 regulates PPARG expression and adipogenesis**.

(A) Validation of the efficiency of FABP5 siRNA in OP9 preadipocytes assessed by carrying out immunocytochemistry. Bar plots show mean +/– SEM from 3 technical replicates with approximately 5000 cells per replicate.

(B-C) Rosiglitazone, a PPARG agonist which binds to the same binding pocket in PPARG as fatty acid, bypasses the need for FABP4 to upregulate PPARG expression during adipogenesis. Knockout of FABP4 and FABP5 impairs adipogenesis in 3T3-FF42A preadipocyte cells induced to differentiate by the standard protocol of adding insulin. The addition of 1 μM Rosiglitazone rescues the loss of adipogenesis in FABP4-KO, FABP5-KO, and DBKO 3T3-F442A cells. Scale bar is 30 μm.

**Figure S2: Increasing FABP4 expression by CRISPRa increases PPARG expression**.

(A) Induction of FABP4 expression 48 hours post transfection in the C3H/10T1/2-CRISPRa-SAM cells using either empty vector (EV) or guide RNA targeting FABP4 promoter region (sgFABP4). The cells were induced to differentiate by addition of the adipogenic cocktail DMI (1 μM dexamethasone, 250 μM IBMX, 1.75 nM insulin) for 48 hours and refreshed the medium with insulin alone for another 48 hours followed by qRT-PCR analysis of FABP4, PPARG and Adiponectin (AdipoQ) expression, data are normalized to 18s. Three biological replicates were used, student T test, 2 tail, type 2 was applied for statistical analysis. Values represent means ± SEM. ***p<0.001, ****p<0.0001.

(B) FABP4 expression was measured 48 hours post transfection using EV or sgFABP4 into the C3H/10T1/2-CRISPRa-SAM cells. The cells were induced to differentiate using the standard 96 hour DMI protocol (see Methods) before being fixed and stained for PPARG protein expression (red), BODIPY (green) and Hoechst (blue). Scale bar: 50 μm.

**Figure S3. Workflow for using CRISPR-mediated genome editing to generate and validate single clones with endogenous PPARG tagged with citrine(YFP) (already existing cells from Bahrami-Nejad et al, 2018) and endogenous FABP4 tagged with mKate2(RFP)**. The different steps in the workflow are described in detail in the Methods section.

**Figure S4. Validation of FABP4-mKate(RFP) OP9 cell clones**. The different steps of the validation procedure are detailed in the Methods section.

**Figure S5. Probability analysis to determine the timing of feedback engagement**.

To determine the timing at which the PPARG dynamics in siRNA knockdown cells start to become diverged from that in negative control cells, we calculated the p-value by Kruskal-Wallis test at each time point. By plotting logarithm of p-value versus time, we found -20 could be a suitable value as the threshold. Then we search for the time point at which the logarithm of p-value become, at the first time, less than -20, and use this very time point (red dashed line, indicated as separated time) as the timing from which the PPARG dynamics between knockdown cells and negative control cells start to get remarkably diverged.

## CONTACT FOR REAGENT AND RESOURCE SHARING

Further information and requests for reagents may be directed to, and will be fulfilled by the corresponding author, Dr. Mary N. Teruel.

## EXPERIMENTAL MODEL DETAILS

OP9 mouse stromal cell line [46]

3T3-F442A mouse preadipocyte cell [47]

Immune-deficient mouse-J:NU (Jackson Labs, Cat #007860)

## METHOD DETAILS

### Cell culture and differentiation

OP9 cells were cultured according to previously published protocols [20,23,37,46]. OP9 cells were cultured in 20% Fetal Bovine Serum (FBS) in growth media consisting of MEM-α (Invitrogen, #12561) and 100 units/mL Penicillin, 100μg/mL Streptomycin, and 292 μg/mL L-glutamate (Invitrogen, # 10378-016). To induce differentiation of OP9 cells using the standard 96-hr DMI protocol, confluent cells were treated with a differentiation medium containing a commonly used DMI (dexamethasone/IBMX/insulin) stimulus to initiate adipogenesis. DMI consists of dexamethasone (dex), a synthetic glucocorticoid; 3-isobutyl-1-methylxanthine (IBMX), an inhibitor of phosphodiesterase that increases cAMP levels; and insulin. Applying the DMI stimulus consisted of replacing the media on the cells with growth media plus 10% FBS, 0.25 mM IBMX (Sigma Cat # 7018), 1 μM dexamethasone (Sigma Cat #D1756), and 1.75 nM insulin (Sigma Cat # I6634) (Stimulus I). Forty-eight hours after initiating differentiation, Stimulus I was removed from the cells and was replaced with Stimulus II consisting of growth media plus 10% FBS and 1.75 nM insulin for another 48 hours. As noted in some experiments, rosiglitazone (Cayman, USA) was added to the media to result in a final concentration of 1 μM.

3T3-F442A preadipocytes were grown and differentiated according to established protocols[30]. Briefly, 3T3-F442A cells were cultured in 10% bovine calf serum in growth media consisting of DMEM, 2 mM L-glutamine, 100 U/ml penicillin, and 100 U/ml streptomycin. Cells were passaged when pre-confluent. For differentiation, preadipocytes were grown to confluency on plates coated with collagen-1 (Advanced BioMatrix, 5005). When confluent, differentiation was induced by changing media to growth media + insulin (5µg/ml) for four days. After day 4, the media was replaced with just growth media. This latter media was replaced every two days on the cells until the cells were fully differentiated, which occurred typically around day 10 after induction of differentiation.

Linoleic acid (LA, Sigma-Aldrich L1012) was first dissolved in 100% ethanol to a concentration of 1M, mixed with 10% fatty acid-free BSA (Sigma-Aldrich A8806) in MEM medium, and then incubated at 37°C for 4 hours to form LA-BSA complexes at a final concentration of 1mM LA. Then the 1 mM mixture of LA-BSA complexes was diluted 10-fold in MEM to a stock concentration of 100 μM. This stock concentration was then further diluted with additional MEM before applying to cells.

### Oil Red O staining

To determine lipid accumulation 3T3-F442A preadipocytes were differentiated for 10 days and stained with Oil Red O. The cells were washed with phosphate-buffered saline (PBS) and fixed using 4% paraformaldehyde (PFA) for 1h. Cells were washed using double-distilled H_2_O and incubated with the filtered Oil Red O solution (Cat #O0625, Sigma) for 1h. Cells were washed twice with ddH_2_O and analyzed by bright-field microscopy.

### *De novo* adipogenesis *in vivo*

To induce *de novo* fat pad formation, preadipocytes were grown to near confluency and resuspended in PBS. 3 × 10^7^ preadipocytes were injected subcutaneously at the sternum of 8-week old male athymic mice (Cat. #007850, Jackson Laboratory) as described previously [48]. After 4 weeks, mice were anesthetized with isoflurane and sacrificed by cervical dislocation. Fat pads derived from the implanted cells were excised and weighed.

### Generation of FABP4-KO OP9 cells and FABP4-KO, FABP5-KO, and DBKO 3T3-422A cells

The CRISPR-Cas9 constructs targeting FABP4 and FABP5 were generated based on a previously described protocol [49]. Briefly, different guide RNAs targeting FABP4 or FABP5 were designed (crispr.mit.edu) and oligos including the targeting sequences were annealed and cloned into pSpCas9n(BB)-2A-GFP (PX461; Addgene #48140), used for targeting FABP4, or pSpCas9(BB)-2A-miRFP670 (Addgene #91854), used for targeting FABP5. Constructs were transfected into 3T3-F442A preadipocytes or OP9 preadipocytes via electroporation using the Amaxa Cell Line Nucleofector Kit V (Lonza, Cologne, Germany). Afterwards cells were grown two days in DMEM, 10% BS and 100 U/ml penicillin/streptomycin for 3T3-F442A or for OP9 MEM with L-glutamine, 20% FBS, and 100 U/ml penicillin/streptomycin. FACS-sorting was performed to select for construct marker (GFP and/or iRFP expression), and cells were seeded in 96-well plates (1 cell/well) to analyze them for FABP4 and/or FABP5 expression. Single cells that did not harbor any FABP4-KO were expanded to obtain WT(control) clones. Successfully transfected cells were sorted for positive cells and assayed for loss of FABP4, FABP5 or both proteins.

### Generation of stable, doxycycline-inducible FABP4 overexpression OP9 cell lines

The Lenti-X Tet-On Advanced lentiviral inducible expression system (Clontech Laboratories, Mountain View, CA) was used to generate a doxycycline-inducible protein expression system in our stable OP9 FABP4-KO cell line. This system needs two elements: 1) the Regulator Vector (pLVX-Tet-On Advanced) and the Response Vector (pLVX-Tight). Our FABP4-KO OP9 cells were co-transduced with pLVX-Tet-On Advanced and either pLVX-Tight-YFP-FABP4(WT), pLVX-Tight-YFP-FABP4 (R126Q) or pLVX-Tight-YFP(control) lentiviruses. Cells were then grown in media containing neomycin and blasticidin S to select clones expressing the two constructs, and FACS was used to further enrich for YFP-positive cells.

To carry out experiments, the stable OP9 FABP4-KO cells with inducible YFP-wild type FABP4, YFP-FABP4 (R126Q) or YFP were plated at a density of 8000 cells per well of a 96-well plate at day -1. Cells were grown in MEM with L-glutamine, 20% FBS, and 100 U/ml penicillin/streptomycin. At day 0, the cells were treated with doxycycline 1ug/ml and adipogenic cocktail in MEM with L-glutamine, 10% FBS, and 100 U/ml penicillin/streptomycin.

### Workflow used to generate mouse OP9 cells with endogenously tagged PPARG and FABP4 (Fig S3)

Previously CRISPR-mediated genome editing had been used to tag the N-terminus of endogenous PPARG in OP9 cells [20]. These cells had also been stably transfected with a H2B-mTurquoise(CFP) as a nuclear marker to facilitate cell tracking. We now used these cells and tagged the C-terminus of FABP4 with mKate2, a bright red fluorescent protein [50].

#### A) Construction of DNA Plasmids

To carry out CRISPR genome editing to generate FABP4-mkate2, we used the “double nickase” system which uses two different guide RNAs that create adjacent and opposing nicks in the DNA at the site of insertion [20,51]. To carry out double-nickase genome-editing, two different targeting sequences (sgRNA) directed to the FABP4 locus were designed as described above and inserted into the guide RNA site of two plasmids, pX335-U6-Chimeric_BBCBh-hSpCas9n (pX335) (Addgene plasmid #42335) encoding the SpCas9 D10A nickase. Oligonucleotide duplexes encoding each desired targeting sequence were ligated into the BbsI cut sites ofpx335. Construction and design of the donor template were carried out based on our previously established protocol [20]. The DNA repair template to promote homology directed repair (HDR)-mediated insertion of the fluorescent protein (mkate2) was constructed by inserting the cDNA of mkate2-3xGly flanked by two 800 bp homology arms into the entry vector backbone pENTR1a (Addgene Plasmid #17398). The pENTR1a backbone vector was digested with EcoRl-HF and BamHl-HF (NEB), and assembled together with three DNA fragments by using Gibson assembly. All constructs were validated by sequencing.

#### B) Transfection

1 mg of each of the two pX335 guide RNA/SpCas9n constructs and 5 mg of the Citrine donor template were transfected into 1 million OP9 cells using Lipofectamine 2000 (Invitrogen) following the manufacturer’s protocol.

#### C) Clone Selection by Single-Cell FACS

Clone selection were carried out based on our previous protocol [20]. In briefly, seven days post-transfection, single cells expressing Citrine were sorted into separate wells of 96-well culture plates and allowed to grow. We chose to wait 7 days post transfection to avoid false positive fluorescent signal originating from the un-integrated donor DNA plasmid.

#### D) Stimulus Response Test

Once the single-cell colonies grew to 50% confluency, each colony was passaged into wells on two different 96-well plates. One plate was used to expand the colonies, and the other half was imaged using a Molecular Devices MicroXL fluorescence imaging system to select for clones with correct localization of the Citrine signal and the appropriate response to stimuli. Before imaging, PPARG/FABP4 clones were stimulated for 24 hours with DMI to induce expression of mCitrine-PPARG /mKate-FABP4.

#### E) Differentiation Capacity Test

Clones that expressed Citrine were further characterized for their differentiation capacity using the standard four-day adipocyte differentiation protocol detailed under ‘‘Cell Culture and Differentiation.’’ Clones that acquired mature adipocyte morphology and accumulated lipid droplets in response to DMI treatment were expanded and subjected to further validation steps.

#### F) Further Validation

Validation of mCitrine-PPARG /mKate-FABP4 Clones. We first performed genomic PCR with different primer sets to look at the genotype and verify correct insertion of mKate into the FABP4 locus. The first PCR was performed using primers annealing to regions flanking the site where mKate is inserted (Table S5; Figure S3A). Using this set of primers, FABP4 tagged clone2 or 4 were shown to be heterozygous, and the PCR products were subjected to sequencing. The second set of PCRs was performed using primers annealing end of the inserted mKate. (Table S6; Figure S3B). Both reactions resulted in a single band, indicating that mKate was correctly inserted into the the FABP4 locus in clone2 or 4. Next, western blot analysis of mKate-FABP4 clone2 was performed using anti-RFP antibody and anti-FABP4 to verify protein expression and to check for the correct molecular weight of the tagged protein (Supp Figs S3C and S3D). The size compatible with the correct predicted molecular weight of mKate-protein fusion were shown for each clone. The anti RFP blot shows the expression of tagged-FABP4 and did not detect any free RFP. Next, Southern blot analysis was performed to confirm locus-specific knock-in using a probe directed toward mKate. All examined clones showed the presence of a specific copy of mKate within the genome, as evidenced by the detection of a single band of expected size (∼2kb) in the Southern blot (Figure S3E). Finally, immunohistochemistry analysis was used to validate co-localization of the mKate fluorescence signal with the immunohistochemistry signal of the FABP4 proteins throughout four days of differentiation. Since it passed all the validation criteria described above, the FABP4-2 clone was used for all the time course measurements in the current manuscript.

### Immunoblotting

For SDS-PAGE, 20 ug protein per lane was used. Proteins were detected with anti-PPARG (1:1000, Santa Cruz Biotechnology, sc-7273), anti-FABP4 (1:1000 R&D Systems, AF1443), anti-FABP5 (1:1000 Cell Signaling, 5174), anti-GFP (1:1000 Abcam, ab290) and anti-tRFP (1:1000 Evrogen EVN-AB233-C100).

### Co-Immunoprecipitation (Co-IP)

Twenty-four hours before the start of the experiment, stable FABP4-KO OP9 cells with either inducible YFP, YFP-WT-FABP4 or YFP-FABP4 (R126Q) were treated with doxycycline (1ug/ml) and in alpha-MEM with L-glutamine, 10% FBS, and 100 U/ml penicillin/streptomycin. At t=0 hours, cells were treated with both linoleic acid (100 μM) and doxycycline (1ug/ml). Co-IP’s on the total cell lysates or on subcellular fractions (nuclear and cytoplasmic) were performed at different timepoints using anti-GFP V_H_H coupled to magnetic agarose beads (GFP-Trap_MA, Chromotek) or, as a control, IgG (Chromotek), following the manufacturer’s instructions. The same amount of protein was used for each IP, and the protein concentration was measured using a BSA kit.

### FABP4 Gain-of-Function in pre-adipocytes via CRISPRa SAM

Murine mesenchymal C3H10T1/2 cells stably expressing the CRISPRa-SAM complex were used for gain of function experiments following a previously established protocol [33]. Briefly, to make the sable cell line, the CRISPRa-SAM components (dCas9-VP64 and MS2-P65-HSF1) were delivered in lentiviruses by plasmid co-transfection of C3H10T1/2 cells. The resulting cell line (C3H10T1/2-CRISPRa-SAM) can be used for activation of endogenous genes via chemical transfection with a single guide RNA (sgRNA)-containing plasmid. C3H10T1/2-CRISPRa-SAM cells were maintained in high glucose-DMEM media containing FBS (10%), Penicillin-streptomycin (1%), and blasticidin (2.5 ug/ml) and hygromycin (200 ug/ml) to ensure a retained dCas9-SaM expression. sgRNA sequences targeting FABP4 promoter regions were designed using the ‘SAM genome engineering online tool’ (http://sam.genomeengineering.org/database/), and annealed and ligated into the backbone vector following the original protocol [52]. Correct insertion was verified by sequencing of plasmids. The sequence used was FABP4 (NM_024406.3) CATACAGGGTCTGGTCATGA. Cell transfections were performed as previously described [33].

For gene expression analysis (RT-qPCR), 300,000 cells were seeded per well into a 12-well plate (Day -3), transfected at confluence (Day -2) using Mirus TransIT-X2 transfection reagent (3ul per well; Mirus Bio LLC, US) and 250 ng plasmid DNA (either empty vector or FABP4 sgRNA). For staining experiments, 2000 cells were seeded per well into a 96-well Costar Plastic plate. The day after, the cells were transfected with a total of 50 ng sgRNA per well using 0.25 μl TransIT-X2 reagent per well. Two days following transfection at Day 0, all cells received either only DMI or DMI and Rosiglitazone (1 uM) to induce adipogenesis. The non-stimulated cells were used as a control. At Day 2, the media was replaced with media containing only insulin. At Day 4, the cells were either fixed for protein staining or lysed for mRNA analysis

### RNA Isolation and Quantitative Polymerase Chain Reaction (qPCR)

Cells were lysed in RLT buffer (β-Me, 1:100) on ice; RNAs were isolated using RNeasy spin columns (Qiagen, CAT. 74106) and cDNA synthesized using the qScript kit (Quantabio, Cat. 101414-098) according to the manufacturer’s instructions. Real-time PCR was performed using the GoTaq qPCR Master Mix (Promega, Cat. M3001) in LightCycler® 480-Roche System according to the supplier’s manual. Raw CT data were normalized to 18S reference gene following the ΔΔ-CT calculation. PCR primer sequences were synthesized by Elim Biopharmaceuticals Inc (USA, CA) and are listed in Supplementary Table S8.

### Fluorescent imaging

Imaging was conducted using an ImageXpress MicroXL (Molecular Devices, USA) with a 10X Plan Apo 0.45 NA objective. Live fluorescent imaging was conducted at 37°C with 5% CO_2_. A camera bin of 2×2 was used for all imaging condition. Cells were plated in optically clear 96-well plates: plastic-bottom Costar plates (#3904) for fixed imaging or Ibidi µ-Plate (#89626) for live imaging. Living cells were imaged in FluoroBrite DMEM media (Invitrogen) with 10% FBS, 1% Penicillin/Streptomycin and insulin to reduce background fluorescence. Images were taken every 12 min in different fluorescent channels: CFP, YFP and/or RFP. Total light exposure time was kept less than 700 ms for each time point. Four, non-overlapping sites in each well were imaged. Cell culture media were changed at least every 48h.

### Imaging data processing

Data processing of fluorescent images was conducted in MATLAB (MathWorks). Unless stated otherwise, fluorescent imaging data were obtained by automated image segmentation, tracking and measurement using the MACKtrack package for MATLAB. Quantification of PPARG- and FABP4-positive cells in fixed samples was based on quantification of mean fluorescence signal over nuclei. Cells were scored as PPARG- and FABP4-positive if the marker expression level was above a preset cut-off determined by the bimodal expression at the end of the experiment.

For live imaging data of OP9 cells, the CFP channel capturing H2B-mTurquoise fluorescence was used for nuclear segmentation and cell tracking. Obtained single-cell traces were filtered to removed incomplete or mistracked traces according to the following criteria: cells absent within 6 hours of the endpoint, cell traces that started more than 4 hours after the first timepoint, and cells that had large increase or decrease in PPARG intensity normalized to the previous timepoint. If cells were binned according to their PPARG expression, cells were binned based on their mean nuclear PPARG expression at the described timepoints. The percent of PPARG high cells was assessed by counting cells that above the PPARG threshold at that time point and dividing by the total number of cells at that time point.

### Estimating a differentiation commitment point (i.e. PPARG threshold)

The timepoint at which cells commit irreversibly to differentiate, or PPARG threshold, is calulcated for each experiment as described in [20,37]. Briefly, PPARG values at the end of a differentiation experiment typically exhibit a bimodal distribution. In order to estimate a commitment point, PPARG values at the last frame of the experiment was fit to a 2 component gaussian mixture model. Cells were then classified as either differentiated or undifferentiated based on whether they more closely associated with the high or low component of the mixture model, respectively. The commitment point was then assessed as the value of PPARG at the 48-hour time point, before the stimuli was removed, that predicted the final differentiation classification with a false positive rate of less that 5%. In experiments where multiple conditions are present, the gaussian mixture model was only fitted to the negative control and the commitment point was selected based on the negative control model and applied to all other conditions in the same experiment.

### Measuring protein decay rates using cycloheximide

Protein decay rates were quantified as previously described [20,27]. Briefly 10,000 OP9 cells were seeded in 96-well plates) one plate for each timepoint. Cells were induced to differentiate with DMI for 24 hours. Cyclohexamide was added to the media at a final concentration of 30 μM. Cells were fixed and stained at different times after addition of cyclohexamide, and immunofluorescence was used to quantify protein concentration. Half-lives were obtained by fitting first order exponential decay curves to the data.

### Statistics

Unless specified otherwise, data are expressed as mean +/- standard error of the mean (S.E.M). Live traces are expressed as median +/- interquartile range (25^th^-75^th^ percentiles). For histograms with a y-axis labeled “Fraction of Cells,” each histogram (not each plot) is normalized to the total number of cells in the population of that histogram such that all bars in the histogram add to 1. Results are representative of at least two independent experiments.

In Fig 4, to determine the timepoint at which the PPARG dynamics in siRNA knockdown versus control cells start to become distinct from each other, we calculated the p-value by Kruskal-Wallis test at each time point. By plotting time versus log(p-value), we found -20 could be a suitable threshold for the p-value. Then we searched for the first time point at which log(p-value) became less than -20 and used this timepoint as the time at which the PPARG between these two populations started to become significantly distinct from each other.

