## Supplemental Material for "Early enforcement of cell identity by a functional component of the terminally differentiated state"

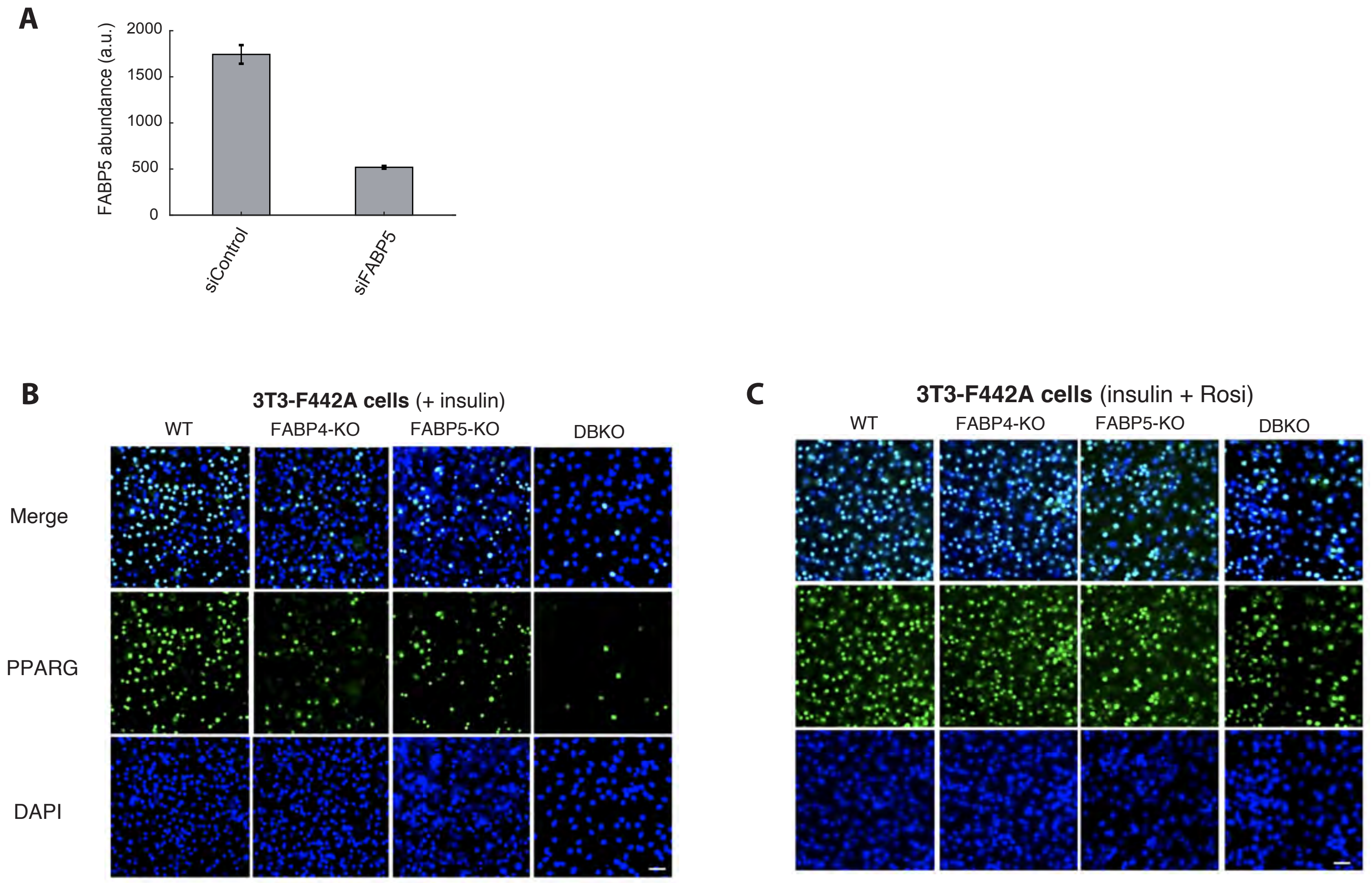


**Figure S1. Additional experiments supporting that FABP4 regulates PPARG expression and adipogenesis.**
(A) Validation of the efficiency of FABP5 siRNA in OP9 preadipocytes assessed by carrying out immunocytochemistry. Bar plots show mean +/– SEM from 3 technical replicates with approximately 5000 cells per replicate.
(B-C) Rosiglitazone, a PPARG agonist which binds to the same binding pocket in PPARG as fatty acid, bypasses the need for FABP4 upregulating PPARG expression during adipogenesis. Knockout of FABP4 and FABP5 impairs adipogenesis in 3T3-FF42A preadipocyte cells induced to differentiate by the standard protocol of adding insulin. The addition of 1 μM Rosiglitazone rescues the loss of adipogenesis in FABP4-KO, FABP5-KO, and DBKO 3T3-F442A cells. Scale bar is 30 μm.

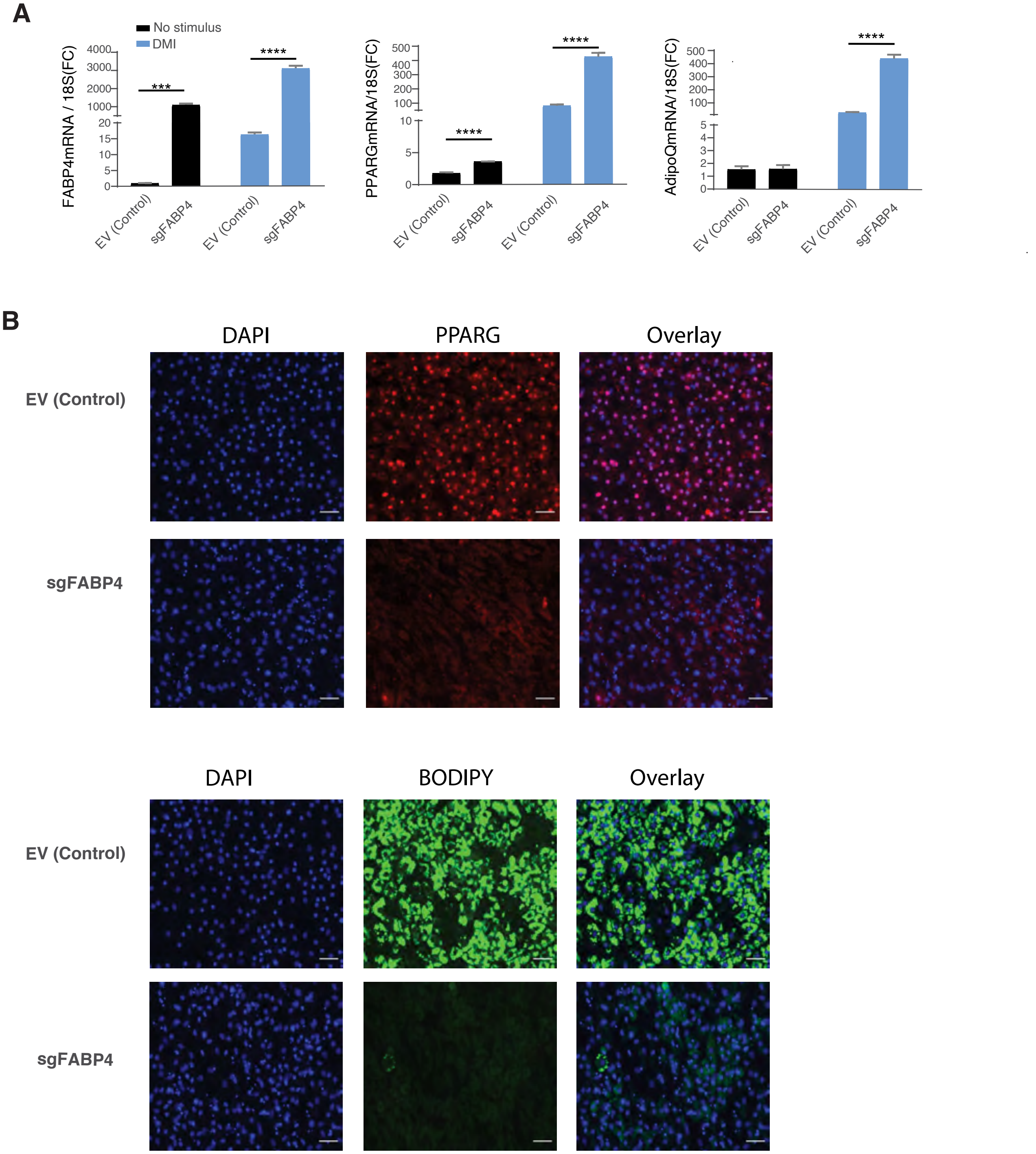


**Figure S2: Increasing FABP4 expression by CRISPRa increases PPARG expression**

(A) Induction of FABP4 expression 48 hours post transfection in the C3H/10T1/2-CRISPRa-SAM cells using either empty vector (EV) or guide RNA targeting FABP4 promoter region (sgFABP4) . The cells were induced to differentiate by addition of the adipogenic cocktail DMI (1 µM dexamethasone, 250 µM IBMX, 1.75 nM insulin) for 48 hours and refreshed the medium with insulin alone for another 48 hours followed by qRT-PCR analysis of FABP4, PPARG and Adiponectin (AdipoQ) expression, data are normalized to18s. Three biological replicates were used, student T test, 2 tail, type 2 was applied for statistical analysis. Values represent means ± SEM. ***p<0.001, ****p<0.0001.

(B) Induction of FABP4 expression 48 hours post transfection in the C3H/10T1/2-CRISPRa-SAM cells using either EV or sgFABP4. The cells were induced to differentiate by addition of the adipogenic cocktail DMI (1 µM dexamethasone, 250 µM IBMX, 1.75 nM insulin) for 48 hours and refreshed the medium with insulin alone for another 48 hours before fixed and stained for PPARG protein expression (red), Bodipy (green) and Hochest (blue). scale bar: 50υM. Adipogenesis assay described in Fig 1C, but now carried out in C3H10T1/2 cells (% PPARG High Cells). Single-cell approach was used to measure expression of FABP4 and BODIPY. Mean FABP4 intensity was obtained by averaging intensities of antibody staining from the nuclei of individual cells. Mean BODIPY intensity was obtained by averaging intensities of BODIPY from the cytosol of individual cells to visualize the lipid accumulation. Bar plots represents mean +/– s.e.m. from 3 technical replicates. Student T test, 2 tail, type 2 was applied for statistical analysis.* p<0.05, ***p<0.001, ****p<0.0001. All data shown are representative of 3 independent experiments.

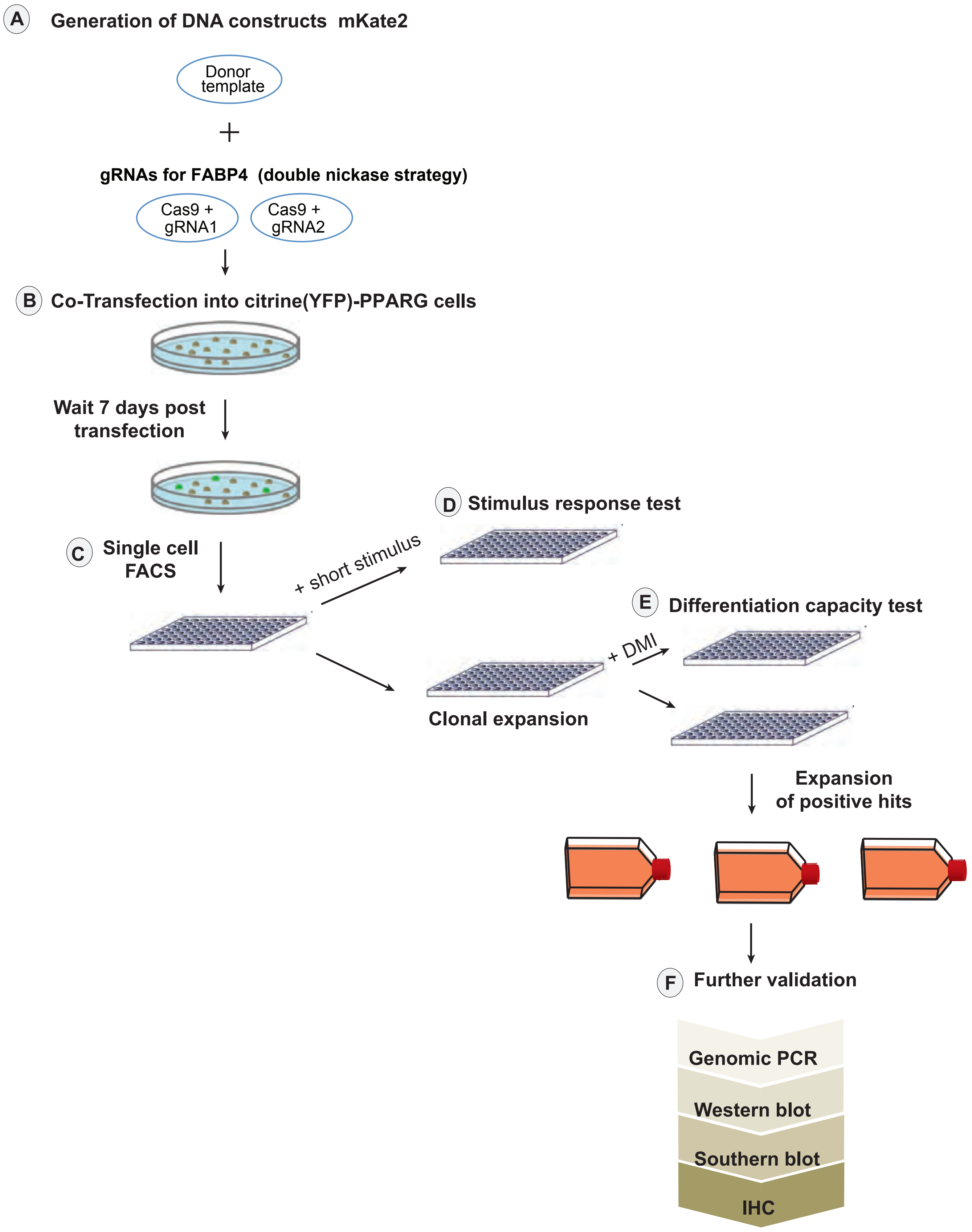


**Figure S3. Workflow for using CRISPR-mediated genome editing to generate and validate single clones with endogeneous PPARG tagged with citrine(YFP) (already existing cells from Bahrami-Nejad et al, 2018) and endogenous FABP4 tagged with mKate2(RFP).** The different steps in the workflow are described in detail in the Methods section.

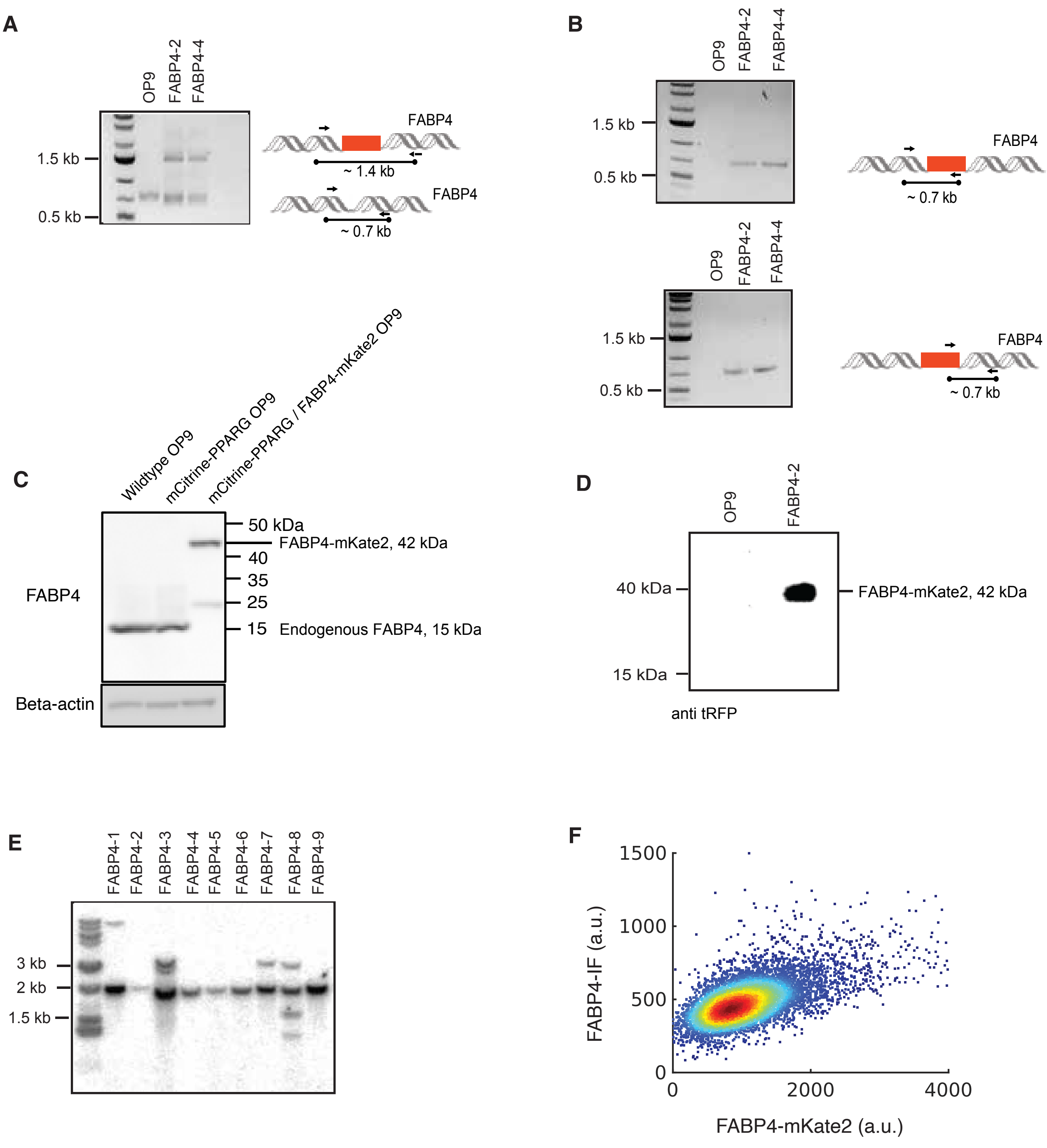


**Figure S4.** Validation of FABP4-mKate(RFP) OP9 cell clones. The different steps of the validation procedure are detailed in the Methods section.

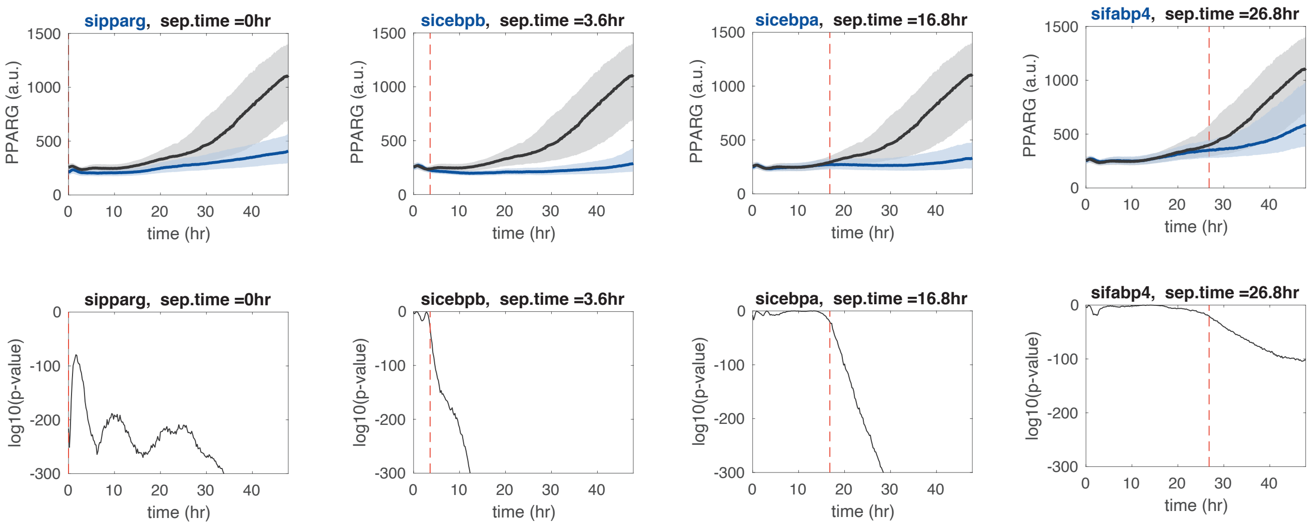


**Figure S5. Probability analysis to determine the timing of feedback engagement.**

### KEY RESOURCES TABLE

| REAGENT or RESOURCE | SOURCE | IDENTIFIER |
|---|---|---|
| **Antibodies** | | |
| Mouse monoclonal anti-PPARgamma (E-8) | Santa Cruz Biotechnology | Cat# sc-7273 |
| Rabbit polyclonal anti-PPARgamma2 | Abcam | Cat# ab45036 |
| Rabbit polyclonal anti-FABP4 | Abcam | Cat# ab13979 |
| Rabbit polyclonal anti-GFP | Abcam | Cat# ab290 |
| Rabbit polyclonal anti-tRFP | Evrogen | Cat# EVN-AB233-C100 |
| GFP antibody for Co-IP | Chromotek | GFP-Trap Magnetic Agarose |
| Mouse monoclonal $\beta$-actin antibody | Santa Cruz Biotechnology | Cat #sc-47778 |
| Rabbit (D1A7T) monoclonal anti-FABP5 | Cell Signaling | Cat #39926 |
| Horseradish peroxidase (HRP)-conjugated anti-mouse | Cell Signaling | Cat #7076 |
| Horseradish peroxidase (HRP)-conjugated anti-rabbit | Cell Signaling | Cat #7074 |
| Goat anti-rabbit IgG (H+L) cross-adsorbed secondary antibody, Alexa Fluor 514 | Invitrogen | Cat #A31558 |
| Goat anti-mouse IgG (H+L) cross-adsorbed secondary antibody, Alexa Fluor 594 | Invitrogen | Cat #A11032 |
| Donkey anti-mouse IgG (H+L) cross-adsorbed secondary antibody, Alexa Fluor 647 | Invitrogen | Cat #A31571 |
| **Chemicals, Peptides, and Recombinant Proteins** | | |
| IBMX | Sigma-Aldrich | Cat #7018 |
| Dexamethasone | Sigma-Aldrich | Cat #D1756 |
| Insulin | Sigma-Aldrich | Cat # I6634 |
| Saponin | Sigma-Aldrich | Cat #47036 |
| Bovine serum albumin | Sigma-Aldrich | Cat #7906 |
| Rosiglitazone | Cayman | Cat #7906 |
| Linoleic acid | Sigma-Aldrich | Cat #L1012-1G |
| BODIPY | Invitrogen | Cat #D-3921 |
| Hoechst 33342 (used like DAPI as a nuclear stain) | ThermoFisher | Cat #H3570 |
| Doxycycline | Sigma-Aldrich | Cat #D3072 |
| **Experimental Models: Cell Lines and Organisms** | | |
| OP9 mouse stromal cell line | Wolins et al., 2006 | N/A |
| 3T3-F442A mouse preadipocyte cell line | Lab of Prof. Emeritus Howard Green, Harvard University | N/A |
| C3H10T1/2 mouse mesenchymal stem cell | ATCC | Clone 8, Cat #CCL226 |
| Immune-deficient mouse-J:NU | Jackson Labs | Cat #007860 |
| **Plasmids** (See Supp. Table 1) | | |
| **Oligonucleotides** (See Supp. Tables 2-8) | | |
| **Cell Lines** (See Supp. Table 9) | | |

| Name | Source | Identifier |
| --- | --- | --- |
| pX335-U6-Chimeric_BB-CBh-hSpCas9n(D10A) | Addgene #42335 | Addgene |
| pSpCas9n(BB)-2A-GFP (PX461) | Addgene #48140 | Addgene |
| tet-pLKO-neo_shFABP4 | This paper | N/A |
| pLVX_tight | Clontech Laboratories Inc | Clontech Laboratories Inc |
| pLVX_tet-on | Clontech Laboratories Inc | Clontech Laboratories Inc |
| pSpCas9(BB)-2A-miRFP670 | Addgene #91854 | Addgene |
| pLVX_tight-FABP4 | This paper | N/A |
| pLVX_tight-FABP4-mutant | This paper | N/A |
| ENTR1A-FABP4-mKate-donor vector | This paper | N/A |

**Table S1: Plasmids**

| **Target** | **Strand** | **Oligonucleotide sequence (5’ to 3’)** |
|---|---|---|
| FABP4_Cterm_1 | Top | *CACCGCATAACACATTCCTAGACAC* |
| FABP4_Cterm_1 | Bottom | *AAACGTGTCTAGGAATGTGTTATGC* |
| FABP4_Cterm_2 | Top | *CACCGTATGAAAGGGCATGAGCCAA* |
| FABP4_Cterm_2 | Bottom | *AAACTTGGCTCATGCCCTTTCATAC* |

**Table S2: Oligonucleotide sequences used to insert sgRNA sequences into the px335.** Guide sequences are targeted to the FABP4 C-terminal. The underlined and italicized nucleotides denote the overhang for ligation of the oligonucleotide duplex into the px335 guide sequence insertion site.

| **Target** | **Strand** | **Oligonucleotide sequence (5’ to 3’)** |
|---|---|---|
| FABP4 | Top | *CACCgGTAATCATCGAAGTTTTCAC* |
| FABP5 | Bottom | *CACCgCCCTTCGAGATCCTTAAGAC* |

**Table S3: Oligonucleotide sequences used for CRISOR KO of FABP4 or FABP5**

| Primer Name | Template | Primer sequence (5' to 3') |
|---|---|---|
| FABP4_homology_region1_FWD | OP9 genomic DNA | AACCAATTCAGTCGACTGCTGTGCC CACAGAGCATCATAAC |
| FABP4_homology_region1_REV | OP9 genomic DNA | TCGCTCACTCCTCCTCCTGCCCTTT CATAAACTCTTGTGGAAGTCAC |
| FABP4_homology_region2_FWD | OP9 genomic DNA | ACTGGGGCACAGATGAGCCAAAGG AAGAGGCCTGGA |
| FABP4_homology_region2_FWD | OP9 genomic DNA | TCGAGTGCGGCCGCGACTCTCTTGA GCATTCAGCCT |
| Fabp4_mKate2_FWD | mKate2 plasmid | TGAAAGGGCAGGAGGAGGAGTGAG CGAGCTGATTAAGGAG |
| Fabp4_mKate2_REV | mKate2 plasmid | CCTCTTCCTTTGGCTCATCTGTGCC CCAGTTTGCTAGGG |

**Table S4: Primers used for PCR amplification of fragments that were joined by Gibson assembly to create donor vectors to insert Citrine at the C-terminal of FABP4 via homologous recombination.**

| **Assay** | **Primer sequence (5' to 3')** | **Amplicon (bp)** |
|---|---|---|
| genotyping FABP4 mKate2 clones | **FWD:** TCTTCCTGGTCTTTGTACCACCCT | **736** (wt allele) **1429**(knock-in allele) |
| | **REV:** CAGGGCAGAAACAAAGCTTCATG | |

**Table S5: Primers used for genomic PCR analysis of the FABP4 CRISPR clones.**

| **Assay** | **Primer sequence (5' to 3')** | **Amplicon (bp)** |
|---|---|---|
| seq. 1 FABP4 mKate2 clones | **FWD:**TCTTCCTGGTCTTTGTACCACCCT | - (wt allele) 712 (knock-in allele) |
| | **REV:** TCCCAGCCGAGTGTTTTCTTCT | |
| seq. 2 FABP4 mKate2 clones | **FWD:**AGAAGAAAACACTCGGCTGGGA | - (wt allele) 739 (knock-in allele) |
| | **REV:** CAGGGCAGAAACAAAGCTTCATG | |

**Table S6: Primers used for genomic PCR analysis to verify the fluorophore integration sites of the FABP4 tagged clones.**

| Primer Name | Primer sequence (5' to 3') |
|---|---|
| mkate2_probe_FWD | CAACCACCACTTCAAGTGCACA |
| mkate2_probe_REV | CTTGAGGTTCTTAGCGGGTTTCTTG |

**Table S7: Primers used for the PCR amplification of a 504 bp probe directed towards Citrine and 505 bp probe directed towards mKate2.**

| Gene | Forward | Reverse |
|---|---|---|
| *Adipoq* | TGT TCC TCT TAA TCC TGC CCA | CCA ACC TGC ACA AGT TCC CTT |
| *Fabp4* | AAG GTG AAG AGC ATC ATA ACC CT | TCA CGC CTT TCA TAA CAC ATT CC |
| *Pparγ2* | TCG CTG ATG CAC TGC CTA TG | GAG AGG TCC ACA GAG CTG ATT |
| *18s* | AGTCCCTGCCCTTTGTACACA | GATCCGAGGGCCTCACTAAAC |

**Table S8. List of primer sequences used for quantitative PCR-based gene expression analysis.**

| Name | Description | Source |
|---|---|---|
| FABP4/PPARG dual tagged OP9 cells | citrine(YFP)-PPARG and mKate2(RFP)-FABP4 cells | This paper |
| FABP4 3T3-F442A Knockout | CRISPR knockout FABP4 in 3T3-F442A cells | This paper |
| FABP5 3T3-F442A Knockout | CRISPR knockout FABP4 in 3T3-F442A cells | This paper |
| FABP4/5 double 3T3-F442A Knockout | CRISPR knockout both FABP5 and FABP4 in 3T3-F442A cells | This paper |
| FABP4KO-OP9 | CRISPR knockout FABP4 in OP9 cells | This paper |
| FABP4KO-YFP-DOX-FABP4 | Doxycycline inducible YFP-FABP4 | This paper |
| FABP4KO-YFP-DOX-FABP4(R126Q) mutant | Doxycycline inducible YFP-FABP4 non fatty acid binding mutant | This paper |
| FABP4KO-YFP-DOX | Doxycycline inducible YFP(control) | This paper |

**Table S9: Cell lines**
